# BGCFlow: Systematic pangenome workflow for the analysis of biosynthetic gene clusters across large genomic datasets

**DOI:** 10.1101/2023.06.14.545018

**Authors:** Matin Nuhamunada, Omkar S. Mohite, Patrick V. Phaneuf, Bernhard O. Palsson, Tilmann Weber

## Abstract

Genome mining is revolutionizing natural products discovery efforts. The rapid increase in available genomes demands comprehensive computational platforms to effectively extract biosynthetic knowledge encoded across bacterial pangenomes. Here, we present BGCFlow, a novel systematic workflow integrating analytics for large-scale genome mining of bacterial pangenomes. BGCFlow incorporates several genome analytics and mining tools grouped into five common stages of analysis such as; i) data selection, ii) functional annotation, iii) phylogenetic analysis, iv) genome mining, and v) comparative analysis. Furthermore, BGCFlow provides easy configuration of different projects, parallel distribution, scheduled job monitoring, an interactive database to visualize tables, exploratory Jupyter notebooks, and customized reports. Here, we demonstrate the application of BGCFlow by investigating the phylogenetic distribution of various biosynthetic gene clusters detected across 42 genomes of the *Saccharopolyspora* genus, known to produce industrially important secondary/specialized metabolites. The BGCFlow-guided analysis predicted more accurate dereplication of BGCs and guided the targeted comparative analysis of selected RiPPs. The scalable, interoperable, adaptable, re-entrant, and reproducible nature of the BGCFlow will provide an effective novel way to extract the biosynthetic knowledge in the ever-growing genomic datasets of biotechnologically relevant bacterial species. BGCFlow is available for downloading at https://github.com/NBChub/bgcflow.

## INTRODUCTION

The number of available genome sequences in public databases has exponentially increased for most bacterial species over the last decade (1). These datasets are becoming crucial for extracting functional knowledge from the diversity found across species pangenomes, and the totality of genes found in available genomes of a species (2, 3). Previous pangenome studies have discovered many biological features common across a particular species, with the first study appearing in 2013 (4–6) and subsequently, specific associated traits to phylogenetic groups within species (4–6). Pangenome analysis has now reached the phylogenetic tree scale (3, 4).

Several pangenome analyses include the investigation of biosynthetic gene clusters (BGCs) involved in producing important natural products or secondary metabolites (7–9). Many known secondary metabolites, which also are referred to as specialized metabolites, have bioactive properties, including antibiotic, antifungal, anticancer, insecticide, food preservatives, and biocontrol, with significant applications in medicinal, agriculture, and food biotechnology (10–12). Advances in genome mining have shown that most bacteria have significant genomic potential that may lead to the discovery of novel specialized/secondary metabolites with diverse bioactivities (13, 14).

This biosynthetic potential varies within and across the species, as demonstrated in prior large-scale genome mining studies (7, 15–20). The pangenome-wide BGC comparisons guide the selection of particular BGCs or strains for further characterization. Several genome mining tools and databases like antiSMASH (21, 22), antiSMASH-DB (23), MiBiG (24), ARTS (25), BiG-SCAPE (26), BiG-SLICE (27), BiG-FAM DB (28) among others are routinely used to predict, search, and comprehensively analyze BGCs. However, each of these tools remains limited based on its scope and the parameters used for accurate predictions of BGC functions or similarity across BGCs. Thus, the integration of various phylogenomic, genome mining, and comparative genomics tools is becoming a central part of investigating secondary metabolite potential and their evolutionary relations across different bacterial genome datasets.

The growth in genome mining tools and available resources poses a strong need for a standardized platform that allows reproducible analyses and easy interoperability of the inputs and outputs required by each genome mining and analysis tool (13). Advances in data science workflows have been transformational in processing sequence data, running standard analytics pipelines, and readily accessing processed output reports (29, 30). For example, different workflow management systems such as Snakemake (31), Nextflow (32), Cromwell (33), and others can assist in job scheduling, logging, and reporting of the key outputs (34). Further, these automated workflow managers can play a critical role in organizing the key results and data in a findable, accessible, interoperable, and reproducible (FAIR) manner assisting researchers in extracting the knowledge from the bacterial genomes efficiently (35). Recently, workflow management systems were applied to efficiently run genome assembly, annotation, or pangenome analysis pipelines (29, 36–38). However, end-to-end large-scale investigation studies for secondary metabolite biosynthetic potential still require significant domain expertise.

Here, we introduce BGCflow, a versatile Snakemake workflow aimed to aid large-scale genome mining studies to comprehensively analyze the secondary metabolite potential of selected bacterial species. BGCflow integrates various genome analytics tools for organizing sample metadata, data selection, functional annotation, genome mining, phylogenetic placement, and comparative genomics. In particular, BGCFlow provides customizable, easy-to-interpret reports, visualizations, and an analytical database for downstream exploratory analysis. Further, we demonstrate the utility of BGCFlow by analyzing publicly available genomes in the genus *Saccharopolyspora*, a rare actinomycete known to be distributed across diverse habitats producing industrially high-value natural products like spinosyns, erythromycins, and others. We show that the BGCFlow-guided interoperable knowledge from different tools and databases can help in the targeted discovery of novel BGCs and more accurate functional clustering of BGCs.

## RESULTS

### Integrated components of BGCFlow present a novel platform to efficiently explore BGC potential across genomes

BGCFlow allows users to collect, organize, explore, and visualize the BGC potential of a large set of genomes either directly collected from the public database (e.g., NCBI (39), PATRIC (40)) or custom-assembled genomes from internal sources. The workflow consists of five different components: i) Snakemake workflow to manage the data files and integrate the execution of the different bioinformatic tools for genome and BGC analysis, ii) project management with efficient configuration of samples and rules, iii) job scheduling and monitoring, iv) analytics database for storing key information and v) Jupyter-based notebooks for reproducible reports, analysis, and visualization (Figure 1). These five components also represent the command line options provided in the BGCFlow wrapper, such as *deploy, init, run, build,* and *serve*.

**Figure 1.**
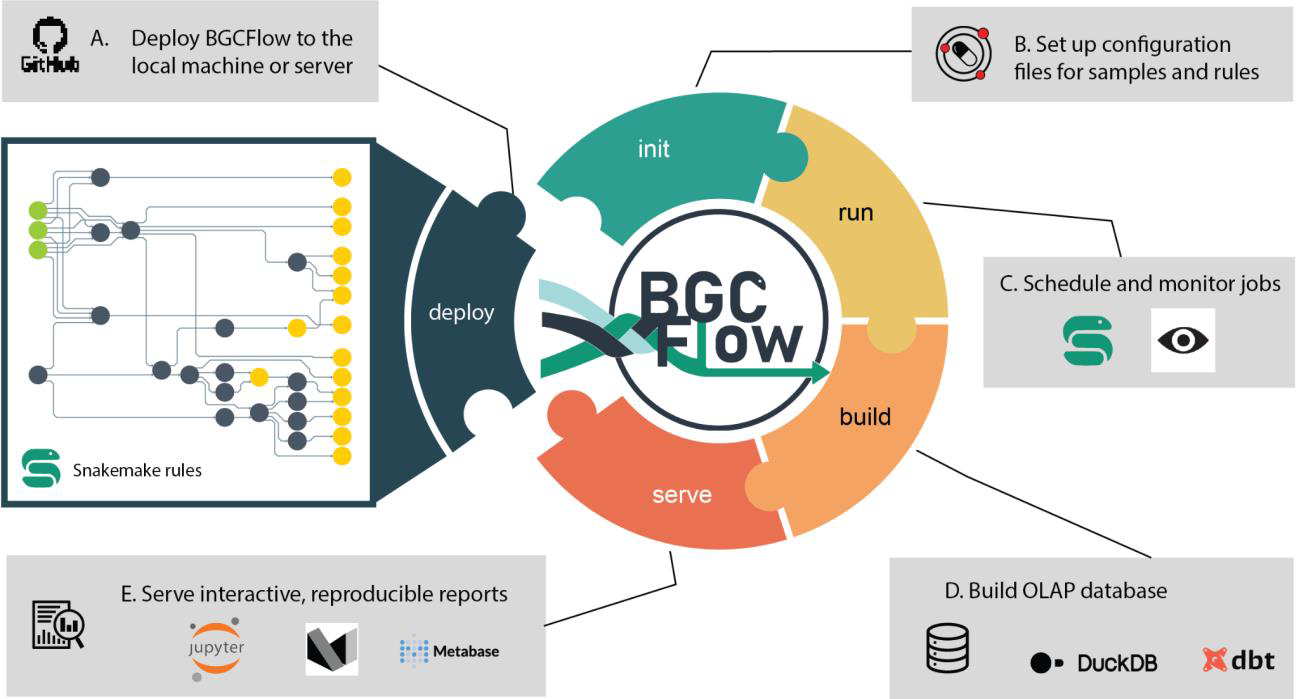
Overview of the BGCFlow components. Overview of the key stages involved in large-scale genome mining analysis supported by BGCFlow commands. (A) Deployment of BGCFlow code along with various rules (command: *bgcflow --deploy*). (B) BGCFlow organizes multiple projects and samples metadata using Portable Encapsulated Projects (PEPs) and a global Snakemake configuration file (.yaml) to select the set of rules to run in a given project (command: *bgcflow --init*). (C) BGCfFlow can automatically run all selected rules using Snakemake job scheduling that can be monitored in live using panoptes (command: *bgcflow --run --wms-monitor*). (D) The generated tables can be readily accessed for downstream analysis through an OLAP database based on DuckDB and the data build tool (DBT) (command: *bgcflow --build*). (E) Processed data is then reorganized per projects, including interactive Jupyter Notebooks and markdown reports (command: *bgcflow --serve*). Additional commands provided in BGCFlow include *--rules, --clone, etc.* Details of different components can be found in Figures S1 to S4.

The main component of BGCFlow includes running a set of well-established genome mining and analysis tools allowing the integration of biosynthetic knowledge provided by different resources. These various existing genome and BGC analysis tools are packaged as configurable Snakemake rules that can select for commonly deployed stages of genome mining studies (Figure 1A). Additional customized rules were added to create seamless interoperability across this various software. In general, the individual rules carry out functions such as input data curation, data wrangling, running external software, managing interoperability between software, and processing the results. BGCFlow contains more than 60 such Snakemake rules (Figure 2B, Table S2), whereas 19 (Table S1) of these can be selected from the configuration to run any of the analyses.

**Figure 2.**
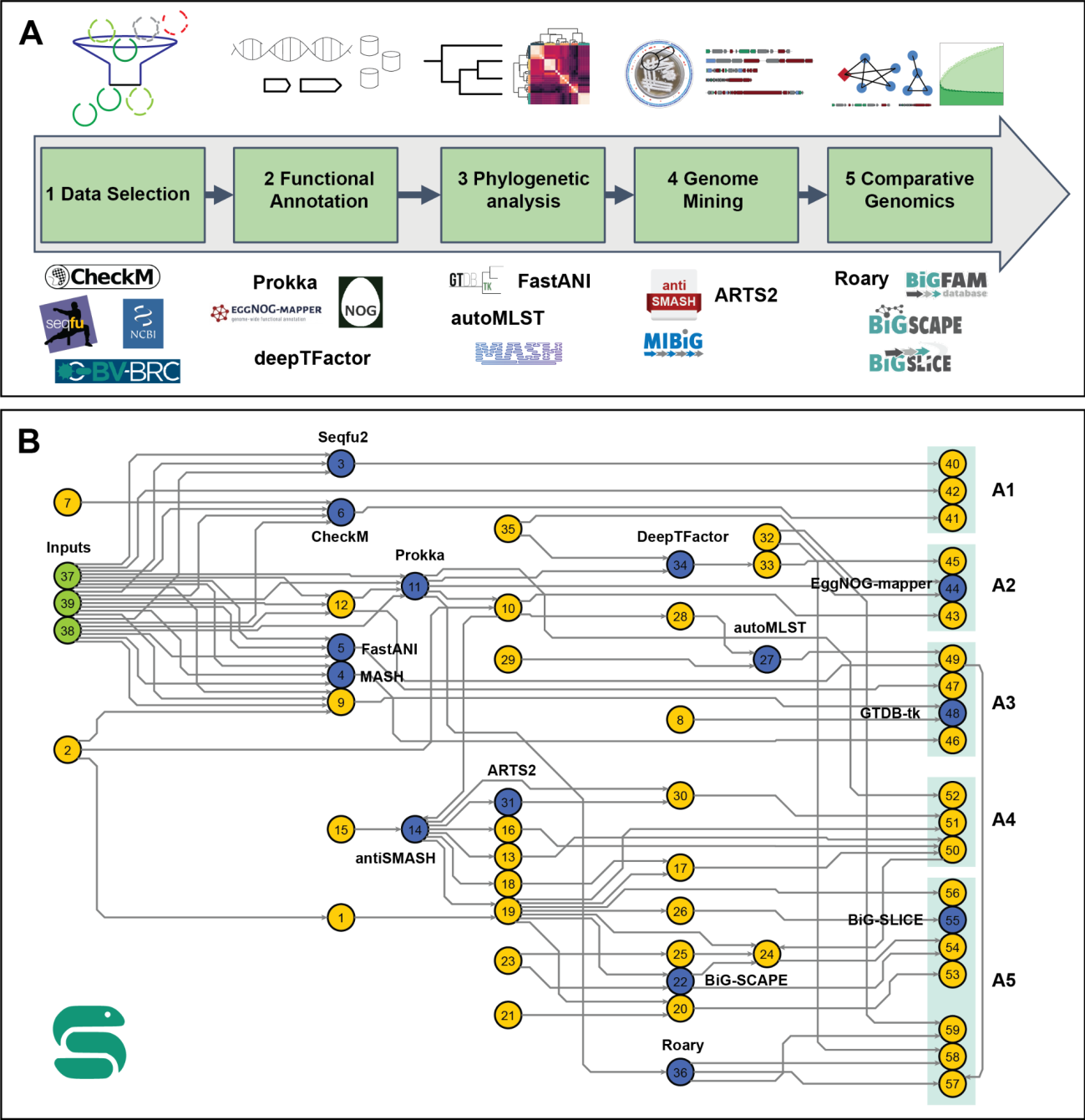
BGCFlow supported rules to carry out common stages of large-scale genome mining studies. A) Overview of the five main analysis stages supported in the BGCFlow. B) Detailed Snakemake rulegraph of BGCFlow. Input nodes are colored green, nodes utilizing main bioinformatic tools are colored blue, and the remaining custom rules for interoperability are colored yellow. Nodes inside green boxes on the right are endpoints for individual pipelines that can be turned on and off by using configuration files. These end-point custom rules can be categorized into the five main analysis types: A1) Data Selection, A2) Functional Annotation, A3) Phylogenomic placement, A4) Genome Mining, and A5) Comparative genomics. Numbers on each node refer to Snakemake rules described in Supplementary Table S1 and S2.

Large-scale studies also include reanalyzing subsets of the initial dataset with different pipelines, leading to multiple projects within a study. For example, when selecting a dataset to build a pangenome, the size of genome samples in the project may change after running the filtering stage based on genome quality or taxonomic placements. To aid in managing sample datasets across multiple projects or studies, we adopted the Portable Encapsulated Project (PEP) specification (41) (Figure 1B, Figure S1). PEP enables users to easily define the project and input configuration via human-readable “.yaml” files for each project and subproject. A typical project configuration includes a path to the table containing a list of sample datasets (e.g., a list of NCBI accession identifiers) as the input, a description of the project, selection of the pipelines to be run by setting the value into TRUE or FALSE. The configuration also allows users to include predefined taxonomic classification results from GTDB-Tk (42) and a list of well-curated GenBank annotations to improve gene annotation while running Prokka (43). BGCFlow can then read multiple PEPs and efficiently assign appropriate computational resources to a selected list of jobs using the Snakemake workflow manager (31). Following Snakemake’s best practices, users can use the dry-run function to get an overview of the number of planned jobs in the current analysis before executing the workflow. The set threads for each rule can be optimized and changed further by users through the Snakemake profile configuration file. Once the Snakemake run is started, BGCFlow monitors the progress across various jobs on a live server using Panoptes (44) (Figure 1C, Figure S2).

A major limitation of many bioinformatics workflows is the lack of interoperability between different tools and analyses. BGCFlow provides an effective solution for investigating the BGCs spread across the dataset by integrating various bioinformatics tools for genomic analyses. After completing all analysis pipelines, the essential data tables are extracted, loaded, and transformed using the data build tool (DBT) (https://www.getdbt.com/) into a database of choice (Figure 1D, Figure S3). By default, BGCFlow utilizes DuckDB as a database. DuckDB is an embedded online analytical processing (OLAP) database suited to handle large volumes of data with complex queries (45). To allow easy interaction with the database, users can leverage Business Intelligence tools, such as Metabase (https://www.metabase.com/), for exploratory data analysis and perform interactive SQL queries. Such a platform provides a user-friendly graphical user interface (GUI) to query and visualize the output for downstream scientific investigation. Finally, BGCFlow provides reproducible Jupyter notebooks for analyzing critical results for each pipeline, enabling customizable data storytelling for users. Using mkdocs (https://www.mkdocs.org/), the standard notebooks are then served as static HTML reports via markdown, inspired by other attempts at reproducible reporting (46, 47) (Figure 1E, Figure S4). With these five components, BGCFlow can significantly assist users in efficiently carrying out large-scale end-to-end exploration of BGCs across bacterial species of interest. In particular, the adaptable and re-entrant nature of the workflow allows users to reenter any stage of the analysis for a selected subset of genomes and tools for more focused exploratory analysis.

### BGCFlow automates common analysis stages of large-scale genome mining studies

A common aim in large-scale genome mining studies is to estimate the biosynthetic potential of strain collection and subsequently prioritize strains or BGCs of interest. Such analysis typically involves various stages, including data selection, functional annotation, genome mining, phylogenetic placement, and comparative analysis. Here, we describe these common analysis stages and their automation using various BGCFlow rules (Figure 2). We note that these stages are not always linear but typically involve cyclic logic for analysis.

#### 1. Data selection based on genome quality and phylogenetic placement

In comparative genome mining analysis, selecting input datasets based on assembly quality and the phylogenetic group is crucial. Many types of BGCs contain repetitive sequence elements, which makes them hard to assemble and causes contig breaks. Thus, genome mining tools such as antiSMASH have the best accuracy with high-quality genomes as an input to capture complete BGCs (21). The data selection stage can be initiated by defining a starting dataset via PEP configuration (Figure S1), which is a “*yaml”* object file pointing to a sample table containing a list of user-provided genome fasta filenames or public NCBI/PATRIC accession identifiers (as “*.csv’’ formatted* table file). Several pipelines can then be run to give an overview of the genome sample quality. The *seqfu* rule of BGCFlow runs Seqfu2 (48) to give a quick overview of assembly statistics such as total genome length, number of contigs, and N50 score. Moreover, the included *checkm* rule’s contamination and completeness results are useful for filtering lower-quality genomes, especially if the study includes metagenome-assembled genomes (49–51). In a few cases, the number of BGCs detected on the contig edge in antiSMASH can also be used as an additional quality filter before running a comparative analysis of BGCs (this would require the *antismash* rule to have been executed).

An appropriate selection of genomes within or across the phylogenetic clades is important in the comparative analysis of BGCs distributed across species or genera. BGCFlow incorporates rules to retrieve taxonomic assignments from GTDB (52) for available genomes in the database. Whereas the GTDB-Tk (42) rule can also assign taxonomy locally for custom genomes. For more specific and customized grouping of input genomes into different phylogroup estimates, we provide the options to calculate MASH (53) and FastANI-based (54) genomic distances across the dataset. A typical PEP for this stage includes a pipeline with the selected rules from *seqfu*, *gtdbtk*, *mash*, and *fastani*. In summary, BGCFlow enables users to iterate over the data selection process using different PEPs with different sample sizes and rules.

#### 2. Consistent functional annotation of the genomes

The consistent gene calling and annotation process across the project ensures that the data are comparable, especially when combining data from multiple sources. Once the input dataset is selected for a particular analysis project, BGCFlow provides a custom annotation rule for Prokka (43) that uses prodigal (55) for gene finding. In particular, the configuration also supports the option for a user-provided list of well-annotated reference genomes of specific species as a priority to annotate while running Prokka. This approach to prioritizing well-annotated genomes is useful for having better species-specific annotations. The GenBank files generated in BGCFlow also add metadata on the time of data generation and the BGCFlow version used.

For extensive functional annotation, BGCFlow can run EggNOG-mapper (56) to predict ortholog groups based on HMM profiles. These extensive annotations include predicted orthologous genes from SEED (57), eggNOG (58), COG (59), GO (60), KEGG (61), and other databases. The classification of genes in COG categories is beneficial for assessing the genomic distribution across biological subsystems. Using the eggNOG rule in combination with Roary (62), which reconstructs the pangenome, provides an overview of the functional content of the entire pangenome of the given species in the project.

The number of functional annotation tools is easily expandable in the BGCFlow framework, as demonstrated by the inclusion of a specialized deep learning-based transcription factor prediction rule for deepTFactor (63). Finally, using the *cblaster* rule, it is also possible to generate a nonredundant DIAMOND database (64) of all gene sequences (nucleotide or protein), which can then be used to query particular genes or gene clusters across the project (65). This analysis stage can include a pipeline with the selected rules such as *prokka*, *eggnog* (or *eggnog-roary*), *deeptfactor* (or *deeptfactor-roary*), and *cblaster-genome*. Thus, BGCFlow enables users to easily access the various functional annotations across all genomes to increase interoperability in the analysis.

#### 3. Phylogenetic analysis

One of the common questions in genome mining studies is to investigate the phylogenetic placement of the BGCs across or within species. As described earlier, the data selection stage also considers the taxonomic placement of genomes using GTDB or custom phylogroup placement using MASH or FastANI (42, 53, 54). A more detailed phylogenetic tree can also be reconstructed using autoMLST based on multi-locus sequence analysis of a few conserved genes with functions (30 by default with a maximum of 100) (66). The autoMLST tree provides a reference for studying the BGC distribution across the project. The template Jupyter Notebook provides basic tree visualization using ggtree (67) and a guide to extract tables for external visualization using iTOL (68). Users can always use the output in Newick format and various BGC metadata tables to visualize the results in other tools. Alternatively, the pangenome rule of running Roary provides a tree based on the alignment of genes in the core genome (62). This analysis stage provides essential data for studying the evolutionary relationships of BGCs across species or genera.

#### 4. Genome mining and overview of BGCs

This pivotal stage of BGCFlow includes rules to identify and analyze the BGCs across the project. More than 70 types of BGCs can be detected using antiSMASH (version 6 or version 7) for detailed investigation (21, 22). In addition to running antiSMASH on all genomes, we also provide summary tables that help get the overview of the BGCs across the dataset assisting in several downstream analyses. With the help of BGCFlow reports, users can also access all of the antiSMASH results interactively on the mkdocs server. BGCFlow configures the *antismash* rule to run ClusterBlast and KnownClusterBlast, allowing users to search for predicted similar BGCs in the antiSMASH-DB and the MIBIG database (23, 69). Targeted genome mining with ARTS2 adds information on potential resistance genes and other ARTS models present in close proximity to the BGCs (25). The presence of particular resistance genes within BGC can help prioritize BGC targets with potential bioactivities such as antibiotics. Another alternative reference that is available is the BiG-FAM database, which contains ∼1.2 million BGCs from public genomes that are grouped into different gene cluster families (GCFs) (28). Using the query BiG-SLICE rule, detected BGCs in the project can be mapped to GCFs in the large BiG-FAM database (27). In general, this stage of BGCFlow allows the integration of the most popular genome mining tools and databases with interoperability between the results.

#### 5. Comparative analysis of BGCs and pangenome

When comparing a large dataset of BGCs, a common approach is to dereplicate BGCs by grouping them into different GCFs, which encode similar secondary metabolites. BGCFlow supports two commonly used tools, BiG-SCAPE (26) and BiG-SLICE (27), to predict GCFs in the given dataset. In addition to the previously explained query feature, the BiG-SLICE also offers fast calculations for GCF clustering in a given dataset and is usually deployed for much larger datasets. The BiG-SCAPE is typically slower but more accurate at identifying GCFs as it uses a combined distance metric based on the organization of protein domains and sequence similarity. In several cases, an optimal scenario is to do rapid GCF clustering using BiG-SLICE followed by BiG-SCAPE on the GCFs subfamilies for the BGCs of interest. The outputs, including similarity networks, BGC, and genome metadata, can directly be imported into the network visualization tools such as Cytoscape, displaying the diversity of GCFs present in the selected dataset.

On top of GCFs calculation using multiple genome mining tools, BGCFlow provides the possibility to integrate information from different genome mining tools and databases like BiG-FAM, ARTS, and MIBIG. Such an integrated approach can guide more accurate GCF definitions and targeted comparative analysis. For example, the sequence similarity network for BiG-SCAPE can be enriched to provide an information graph where detected BGCs, MIBIG entries, BiG-FAM GCF models, and ARTS model are inserted as nodes, and the edges represent the connections across these resources. By following the provided guidelines, users can further expand this network with other connections.

In addition, the cblaster (65) rule provides a database of all genes in BGCs in the project, where users can then run a BLAST search of multiple profiles to find BGCs or genes of interest. This is a handy approach for BGC prioritization by querying for BGCs with specific profiles, such as PFAMs or SMCoGs encoding for special enzymes that support the modification of secondary metabolites. Finally, studying BGCs in the pangenome context can reveal interesting features of secondary metabolites (7, 9, 70). BGCFlow can reconstruct the pangenome of the genomes in the project (usually for a particular species) using Roary, which clusters similar protein ORFs as orthologous groups and provides basic pangenome characteristics, including gene presence-absence matrix (62). As explained in the functional annotation stage, a combined EggNOG and Roary rule assigns the COG category and other annotations to all genes detected in the pangenome. The approaches in this stage are valuable for analyzing GCF distribution and the pangenome of the dataset, providing a guide for further prioritization of strains or BGCs. Further, we provide sub-workflows within BGCFlow that can take BGC samples as inputs to run this stage of analysis on a selected set of BGCs.

A particular study can combine some of these stages by selecting various rules in the PEP definition, where the results can be integrated using DuckDB or Jupyter Notebooks for exploratory analysis. In the next section, we will demonstrate how these stages can be easily carried out using BGCFlow for selected genomes of the *Saccharopolyspora* genus.

### BGCFlow guided data quality selection and phylogenetic analysis of Saccharopolyspora spp

As a case study, we used BGCFlow to investigate the phylogenetic distribution of BGCs across *Saccharopolyspora spp.* Actinomycetes are an important family of bacteria composed of several well-known secondary metabolite producers. Several large-scale studies have revealed the diverse biosynthetic potential of various genera like *Streptomyces* (70, 71), *Amycolatopsis* (17), *Salinispora* (72), and others. In this study, we selected the *Saccharopolyspora* genus to demonstrate BGCFlow because of the following: i) many secondary metabolites like erythromycin, spinosyn, and others produced by *Saccharopolyspora* strains are of great importance to the medicinal (erythromycin), agriculture (spinosyn), and food industry, for example in Chinese liquors (73–77), ii) there are no comprehensive studies on genome mining across this underexplored genus (73, 78), iii) the size of the dataset was sufficiently large for demonstrating all the steps of the BGCFlow.

*Saccharopolyspora* is a multi-spored actinomycete genus that was first isolated from sugar cane bagasse in 1975 (79). Over the years, *Saccharopolyspora* species have been isolated from various natural habitats like soil, desert, fodder, marine sponges and tunicates, deep-sea sediments, salt lake, and stony corals (73). Currently, as many as 46 species (https://lpsn.dsmz.de/genus/saccharopolyspora) and up to 74 strains are reported in the DSMZ databases (80). In this study, we selected 42 genomes from the NCBI RefSeq database on 25 October 2022 belonging to the *Saccharopolyspora* genus with all assembly qualities (Data S1). The number of *Saccharopolyspora* genomes has increased significantly from 3 in 2012, 13 in 2017, to 42 in 2022 (Figure S5). In the first stage of data curation and selection, we created a PEP with rules checkm, gtdbtk, and seqfu to access the information on the quality of the genome assemblies and phylogenetic classification (Data S1). Here, the 42 genomes were classified into three categories based on N50 and number of contigs: a) 16 genomes of high-quality (N50> 5 Mbp), b) 10 genomes of medium-quality (number of contigs < 50), and c) 16 genomes of low-quality (number of contigs >= 50) (Figure S5). The CheckM analysis showed that most assemblies are over 99% complete and under 4% contaminated (Figure S5) (50). The GTDB-based taxonomic placement led to the reclassification of 42 genomes into more than 22 species belonging to three genera: *Sacharopolyspora* (27 genomes), *Sacharopolyspora_C* (8 genomes), and *Sacharopolyspora_D* (7 genomes) (Figure S5)(52). The species assignment from NCBI to GTDB was updated for a few species; for example, *S. endophytica*, *S. terrae*, and *S. karakumensis* were reclassified into one GTDB-species *S. endophytica*. A previous study based on 16S rRNA genes assigned these three as separate species while noting that they belong to a similar cluster (78). Reclassification of taxonomy based on GTDB provides a more structured basis for downstream comparative analysis.

For the next PEP example, we chose the subset of 26 genomes with medium to high-quality genomes (Data S2). As the number of species defined using GTDB was still higher for effective comparison, we used the MASH-based hierarchical clustermap to group the genomes into eight phylogenetic groups (P1 to P8) with the lowest k-means distance (Figure S6). These phylogroups represented optimal clustering at the level between genus and species and were used in downstream visualizations. The genomic properties such as genome length, GC content, and number of BGCs were associated with the defined phylogroups (Figure 3). The genome length varied from 5.8 Mbp to 9.6 Mbp with an average of 7.6 Mbp, whereas the GC content varied from 68% to 73% with an average of 70%. The phylogroup P7 with four genomes of *S. spinosa* and *S. pogona* had the largest genomes and lowest GC content, along with a high number of BGCs. Similarly, the phylogroup P6 of *S. erythrea* also had larger genomes with a high number of BGCs and higher GC content. *Saccharopolyspora hordei* A54 isolated from fodder (81) had the minimum number of 13 BGCs in its ∼5.78 Mbp-sized genome. Similarly, *Saccharopolyspora rhizosphaerae* H219 isolated from rhizosphere soil (82) had only 14 BGCs in its ∼5.82 Mbp-sized genome. A more detailed phylogenetic tree was also constructed using autoMLST based on 30 common genes, which generally matches with the phylogroups as well. The number of BGCs across this phylogenetic tree was quite diverse (Figure 3). Thus, the selected dataset of 26 genomes and the phylogenetic tree were used in more specific comparative analysis in the subsequent stages.

**Figure 3.**
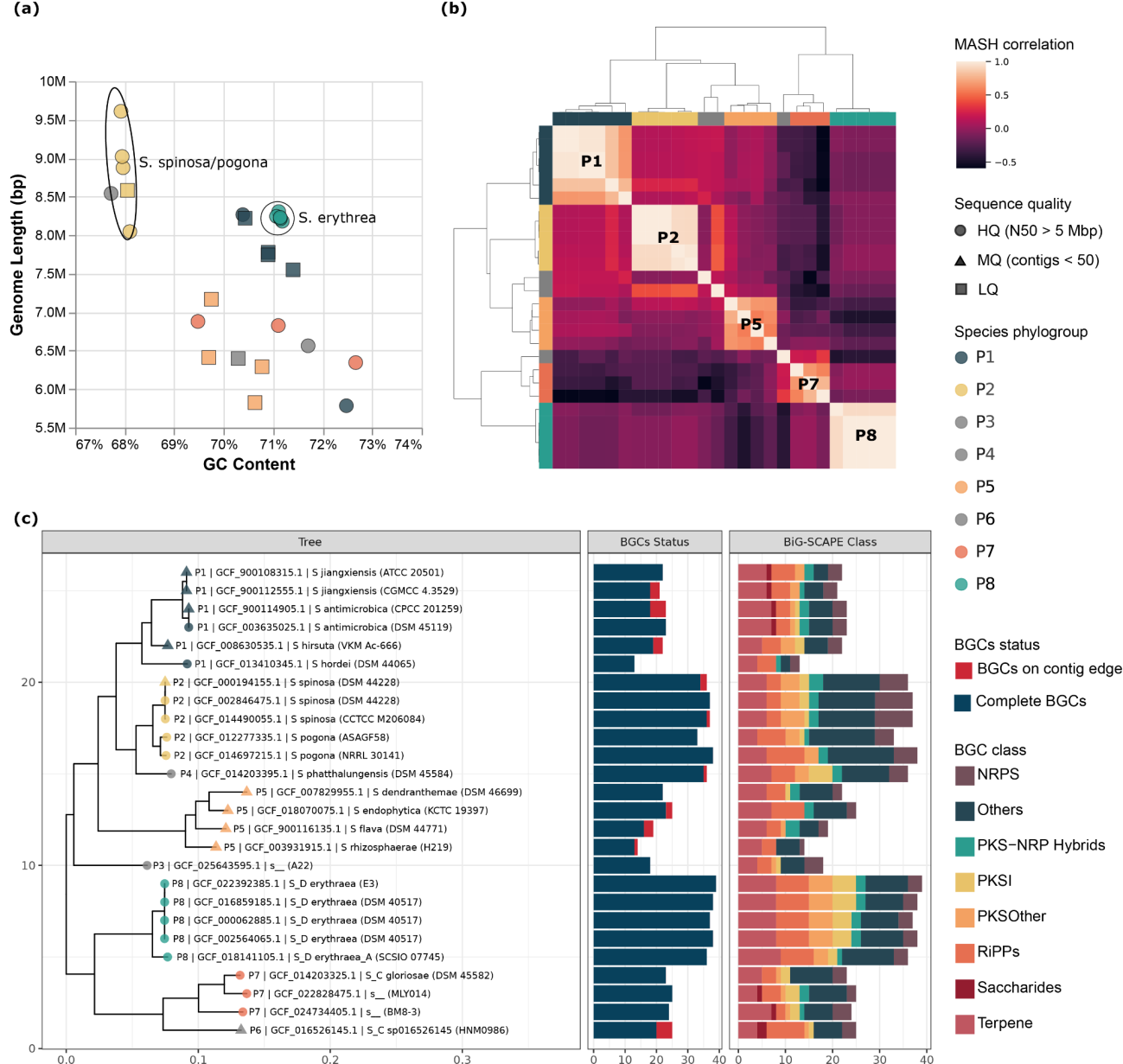
Overview of genomes and detected BGCs in the *Saccharopolyspora* genus A) Scatter plot representing the distribution of genome lengths against GC content across the dataset of 42 genomes of all quality, where the colors represent phylogroup and shapes represent sequence quality (Data S1). B) Clustermap based on MASH-based genomic similarity classifying 26 genomes into eight phylogroups of species (P3, P4, P6 with one strain each) (Data S2). C) Phylogenetic tree calculated using autoMLST displaying the distribution of the number of BGCs. Tree node colors display MASH-based phylogroup and shapes represent the sequence quality. The first stacked barplot represents the number of BGCs on the contig edge or complete. The second stacked bar represents BGCs of different classes as defined in the BiG-SCAPE.

### Integrated genome mining analysis reveals the biosynthetic potential of Saccharopolyspora spp

Here, various genome mining tools were used to predict BGCs and their association with databases such as MIBIG, BiGFAM-DB, and ARTS. The predicted mappings of BGCs against these different databases were used to reconstruct a treemap distribution (Figure 4a). Using antiSMASH, a total of 724 BGCs were predicted across 26 Saccharopolyspora genomes with a median of 25 BGCs per genome (Data S2). The BGCs were distributed across various types such as Saccharides (8 BGCs), PKS-NRP_Hybrids (37 BGCs), PKSI (45 BGCs), PKSother (60 BGCs), NRPS (82 BGCs), RiPPs (90 BGCs), Terpene (159 BGCs), and Others (243 BGCs). Based on the antiSMASH KnownClusterBlast similarity of greater than 80%, a total of 122 BGCs were mapped to the 16 MIBIG database entries of characterized secondary metabolites. Most common hits were geosmin (31 BGCs), ectoine (26 BGCs), 2-methylisoborneol (15 BGCs), and erythreapeptin (14 BGCs) which were found across multiple species. Whereas erythromycin (5 BGCs), flaviolin (5 BGCs), erythrochelin (5 BGCs), spinosyn (5 BGCs), and E-837 furanone (4 BGCs) were detected in specific species.

**Figure 4.**
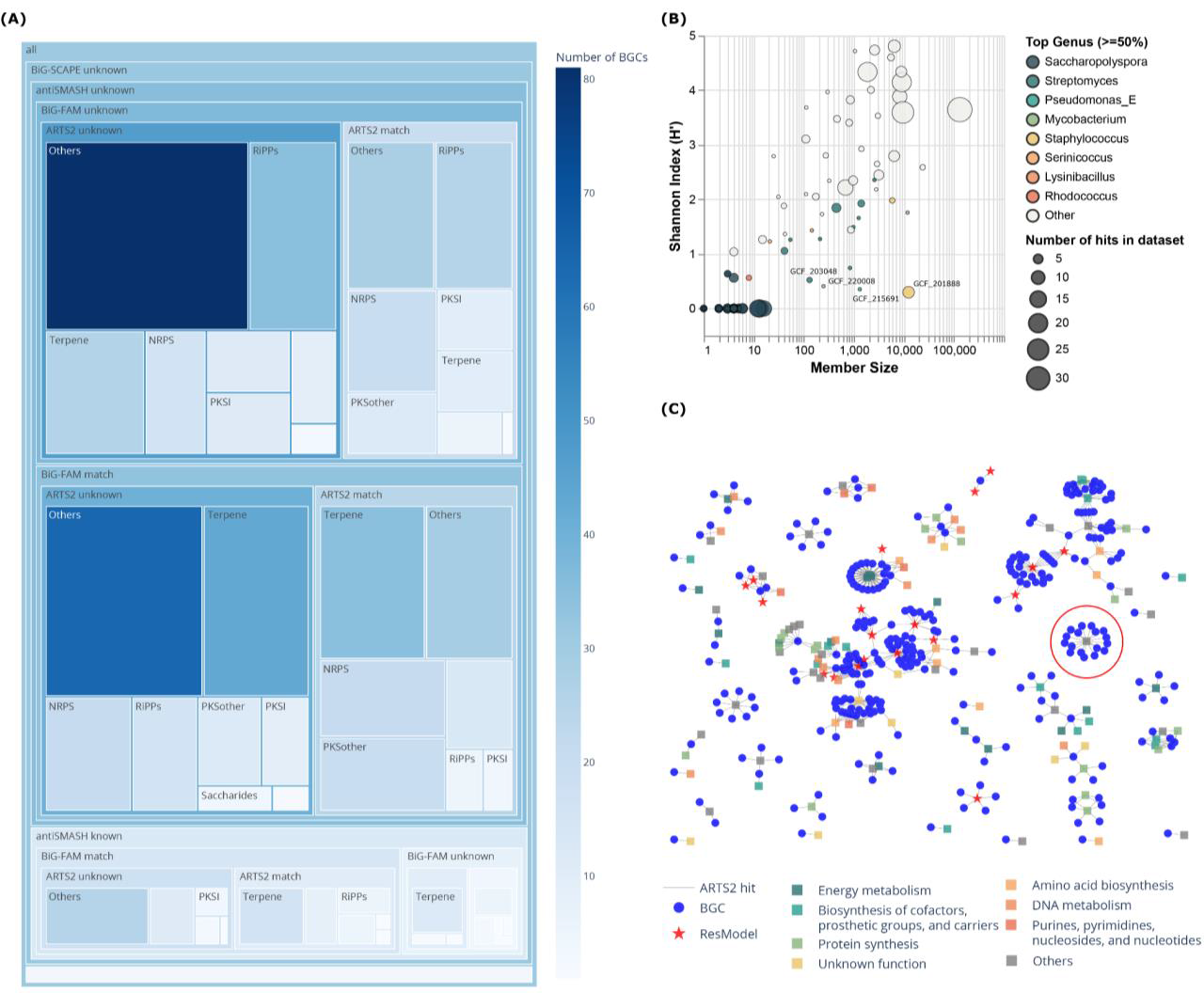
Integrating external BGC knowledgebases of MIBIG-DB, BiG-FAM DB, and ARTS models A) Treemap distribution of the mapping of detected BGCs across different databases such as MIBIG, BiGFAM, and ARTS. BGCs are grouped in antiSMASH known or unknown categories based on greater than 80% similarity to MIBIG DB using KnownClusterBlast function of antiSMASH. Using BiG-SCAPE, BGCs are grouped in known or unknown if the GCF contains MIBIG BGC with a 0.3 cutoff on a similarity metric. Using BiG-SLICE, BGCs have detected a match against BiG-FAM DB of over 1.2 M BGCs detected across public genomes. Finally, the ARTS match represents whether a BGC contains resistance-related genes that can help in prioritization. B) Scatter plot representing the Shannon index against the size of the 130 GCF models detected in the BiG-FAM database. GCFs are given color if there are genera that comprise 50% or more of the total members in that model. C) BGCs with matches against genes from ARTS. Blue color represents detected BGC whereas other colors represent functional classes of genes from ARTS.Highlighted in red circle were the 17 BGCs in proximity to ARTS profile TIGR03997.

The comparison of BGCs against BiG-FAM-DB can be used to investigate if detected BGCs are widely spread across other genomes from the public datasets. The BiG-SLICE-based query resulted in 391 BGCs having similarities against 131 GCF models from the BiGFAM database (Data S3). From the 131 detected GCFs, there were 72 BiG-FAM GCFs that are specifically distributed in the genus *Saccharopolyspora*. Whereas several other GCFs included BGCs from different genera such as *Streptomyces* (47 GCFs), *Amycolatopsis* (31 GCFs), *Kitasatospora* (29 GCFs), *Nocardia* (25 GCFs), *Pseudomonas_E* (22 GCFs), and many others. We further calculated the Shannon diversity index (H) for each of the GCFs representing the distribution of BGCs across different genera (Figure 4B). Of the 59 GCFs with positive Shannon index, 33 were highly distributed across many genera with a Shannon index of greater than 2. The GCF *GCF_201888* with the lowest positive Shannon index (∼0.3) contained 12,444 BGCs distributed across 43 genera with the majority belonging to *Staphylococcus* (∼94.2%) followed by *Acinetobacter* (∼4.2%). The known BGC in this BiG-FAM GCF coded for the biosynthesis of staphylobactin (also known as staphyloferrin B), which is a siderophore with a role in the virulence of *Staphylococcus aureus* (83, 84). The detailed comparative analysis of the predicted BGCs against the MIBIG entry showed that the *Saccharopolypora* genomes indeed possess a BGC that is very similar to the staphylobactin BGC from *Staphylococcus aureus* (Figure S7, Data S4). Some of the other GCFs like *GCF_215691* (Shannon index: 0.35)*, GCF_220008* (Shannon index: 0.41), and *GCF_203048* (Shannon index: 0.52) were found predominantly in genera such as *Pseudomonas_E* (1241 genomes), *Mycobacterium* (227 genomes), and *Streptomyces* (114 genomes), respectively (Figure 4B). These examples particularly highlight the BGCs that are potentially transferred across different phylogenetic groups through horizontal gene transfer events.

Many BGCs coding for known antibiotics also contain genes assisting in self-resistance. By looking for these resistance models, ARTS2 assists in prioritizing novel target screening with potential bioactivity. We detected 296 BGCs that had hits against 599 (130 unique) gene profiles from the ARTS model (Figure 4C, Data S3). The interaction network represented an overview of BGCs in proximity to the ARTS resistance gene models (19 unique) or the core genes models (111 unique) from different functional categories (Figure 4C). We found that 102 BGCs had hits against resistance genes and are more likely to have an antibiotic potential. We also noted that 64 of the 102 BGCs had no similarity to known clusters (either from antiSMASH KnownClusterBlast or BiG-SCAPE results), thus representing underexplored biosynthetic potential of the genus. This analysis further motivated an exploratory analysis of a set of BGCs of unknown function that shared proximity to the same ARTS gene model (17 BGCs), as represented in the last section of mycofactocin-like BGCs of this study.

### Enrichment of the BGC network with knowledgebases revealed more accurate detection and dereplication of GCFs

In order to investigate the diverse biosynthetic potential, it is common to generate sequence-based similarity networks of BGCs that help in identifying distinct Gene Cluster Families (GCFs). Clustering of the detected BGCs in *Saccharopolyspora* using BIG-SCAPE (0.3 cutoff on distance metric) resulted in 330 GCFs. Among these GCFs, only 5 contained known BGCs from the MIBIG database: erythromycin, spinosyn, antimycin, 2-methylisoborneol, and erythreapeptin (Figure 5A, 5C). It is challenging to select an optimal cutoff for defining the GCFs, and thus we evaluated 3 different cutoffs of 0.3, 0.4, and 0.5, respectively (Figure S9). For this study, the selected cutoff is set to 0.3, which is conservative enough to avoid misleading connections with unrelated BGCs and MIBIG references, as demonstrated in less stringent cutoffs (Figure S9). However, we missed many of the known cluster hits due to reasons such as some clusters being hybrid types or having unclear boundaries.

**Figure 5.**
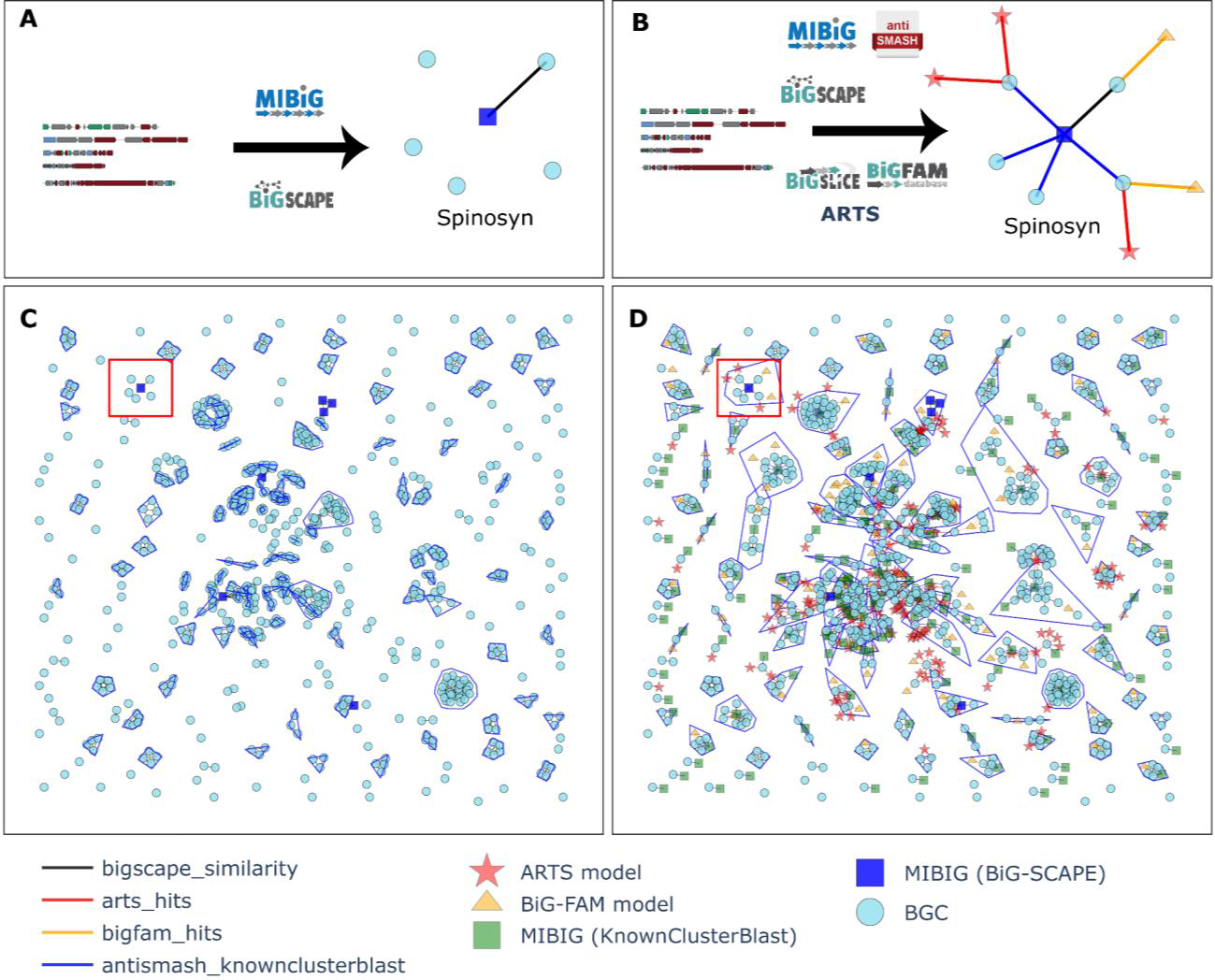
Enriched similarity network with additional connections to different BGC knowledge bases A) Example of BGC similarity network using BiG-SCAPE 0.30 cutoff. B) Examples of the same BGCs with additional connections made using known cluster blast predictions, BiG-SLICE query, and ARTS against the databases of MIBIG, BiGFAM-DB, and ARTS model, respectively. C) Overview of the similarity network of all 722 BGCs resulting in 330 connected components with 19 BGCs assigned to 5 known MIBIG entries and 209 singletons (Data S2). D) Expanded network with information from antiSMASH KnownClusterBlast and BiG-SLICE similarity to BiG-FAM models resulting in 206 connected components with 45 BGCs assigned to 5 MIBIG entries, 89 BGCs assigned to 11 MIBIG entries with >= 80% similarity, and 51 singletons (Data S3). BiG-FAM models 202087, 200946, 210179, 213140, 201682, 201608, 205957, 215277, 201830, and 202082 were removed because the assigned top genus in the model is below 30%.

In this study, we leverage points of reference from available knowledgebases and tools to improve the dereplication accuracy of the GCFs. Our approach proposed within BGCFlow involved generating an extended graph where the BGCs were connected with additional nodes from external knowledgebases such as ARTS (gene profiles), antiSMASH KnownClusterBlast hits (MIBIG BGCs), and BiGFAM-DB (GCF models). The expanded network with information from antiSMASH KnownClusterBlast and BiG-SLICE similarity to BiG-FAM models was used to define more accurate GCFs. The ARTS gene profiles were not used to define the GCFs but can be used to navigate and prioritize GCFs with potentially similar bioactivities. Additionally, the different layers of knowledgebases enable a better explanation of the reason behind the clustering of GCFs.

The enriched network narrows down the 330 GCFs from BiG-SCAPE into 202 connected components with only 50 singletons (previously 209) (Data S3). Using KnownClusterBlast *(>80% similarity)* and matches to BiG-FAM models, we can assign 134 BGCs to 16 MIBIG entries, covering all known *Saccharopolyspora* BGCs and common BGCs such as geosmin and hopene. This information was missed using only the BiG-SCAPE network. One example of using this approach is highlighted by capturing the spinosyn GCF more accurately due to the connections from antiSMASH KnownClusterBlast hits, which were missed in BiG-SCAPE due to the variable boundary of hybrid clusters (Figure 5B, Figure S9, Data S5). Another example included a more accurate definition and a deeper investigation erythreapeptin-like GCF, which was spread across 6 different BiG-SCAPE GCFs (Figure 6C). Overall, the enriched network can be used as a starting point for in-depth comparative analysis within specific GCFs as demonstrated in the next sections, where BGC comparison tools are used on these smaller PEP projects of each GCFs.

**Figure 6.**
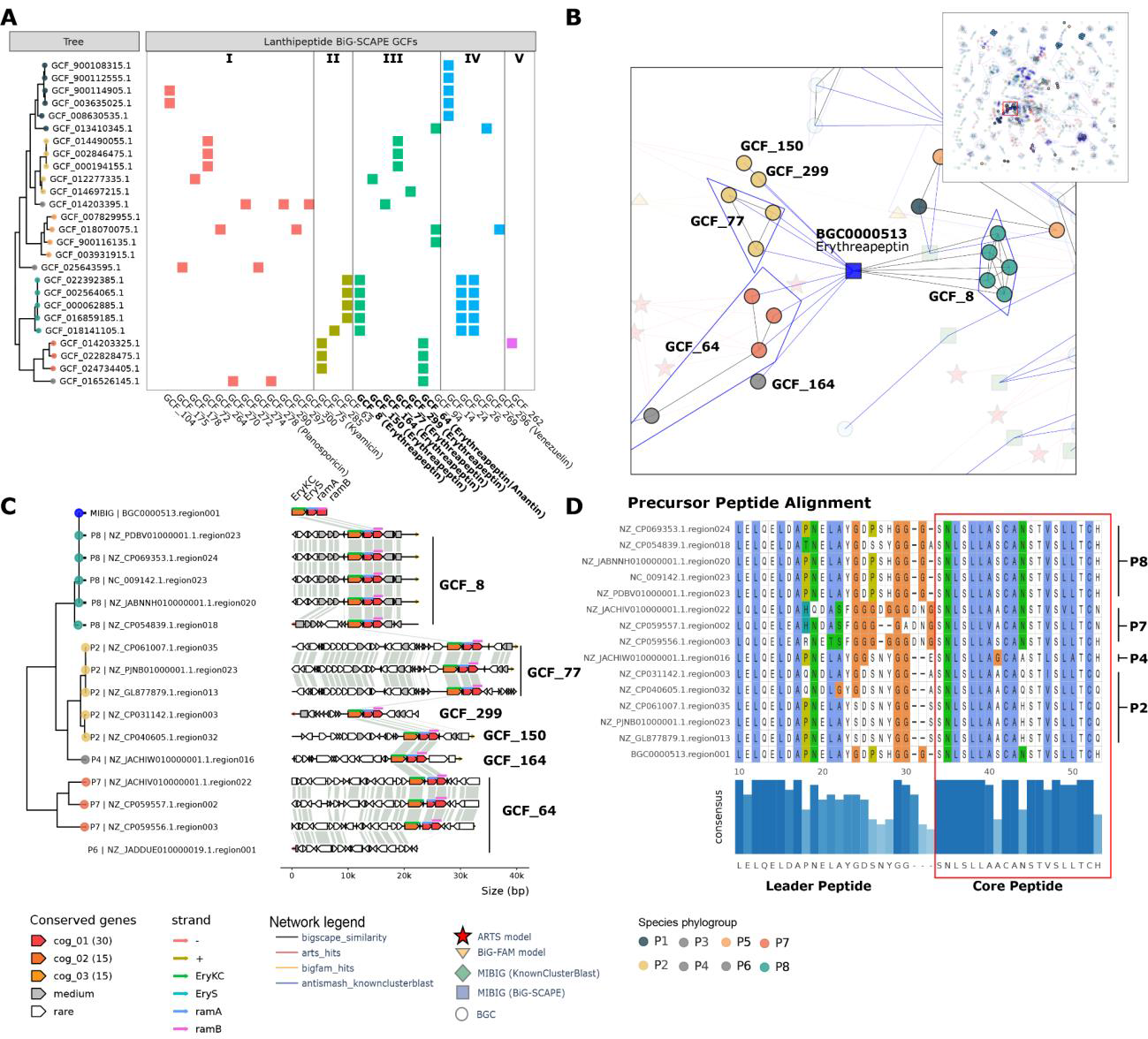
Alignment of lanthipeptide BGCs similar to erythreapeptin A) Phylogenetic tree and distribution of GCFs of all lanthipeptide classes. B) Connected graph with nodes from KnownClusterBlast, ARTS, BiG-FAM. Difference GCFs are highlighted with the same background colors as panel A. C) Alignment of detected erythreapeptin-like BGCs using clinker, ordered phylogenetically (Data S6). D) Sequence variation in precursor peptides highlighting novel variants of erythreapeptin-like lanthipeptides in different phylogroups.

### Distribution and sequence level variation across lanthipeptide GCFs

As an example of in-depth comparative BGCs analysis, we investigated the distribution of BGCs belonging to different types of lanthipeptides. Lanthipeptides consist of five diverse classes of ribosomally synthesized and post-translationally modified peptides (RiPPs) (85, 86). Lanthipeptides are typically represented by peptides containing lanthionine bridges, which are post-translationally synthesized by linking the Cys thiol- and a dehydrated Ser hydroxy group. From the 121 RiPPs detected in the *Saccharopolyspora* genome dataset, we identified 58 BGCs of the various lanthipeptide classes namely class I (15 BGCs), class II (8 BGCs), class III (17 BGCs), class IV (17 BGCs), and class V (1 BGC) (Figure 6A). We found that different classes of lanthipeptides were predominant in specific phylogroups. For example, class II lanthipeptides were detected only in phylogroups P7 and P8. On the other hand class I lanthipeptides were detected in all the other phylogroups except P7 and P8. The class IV lanthipeptides were predominant in phylogroups P1 and P8. The class III lanthipeptides were present in most but phylogroup P1 and P3. Interestingly, these class III lanthipeptides were spread across 6 different GCFs as per BiG-SCAPE but were detected as a single connected GCF based on the enriched network due to KnownClusterBlast similarity (Figure 6B).

We further carried out an in-depth exploratory analysis of the class III lanthipeptide GCF by defining a new PEP with BGC snakefile that used the BGCs of this GCF as samples and BGC alignment tools such as clinker (Data S6). We noted that most BGCs of this GCF contained all of the known biosynthetic genes for erythreapeptin (Figure 6C), a known lantibiotic produced by *Saccharapolyspora* (87). The only exception was a BGC in the phylogroup P6 which had the additional genes but was missing the core biosynthetic genes. We note that the additional genes were different in different phylogroups and thus explain the different GCFs detected using BiG-SCAPE. Finally, we also compared the amino acid sequence variations in the leader and the core peptides of the core gene *eryS* (Figure 6D). We note that the core peptide sequence is generally conserved across all phylogroups with some variations at specific positions. For example, the AA in the position 11 of the core peptide was N in phylogroups P7 and P8, A in P4, and Q or H in P2. We also note that the precursor peptide sequence also displayed variations across different phylogroups. Further experimental studies are required to assess if these variations will lead to lantibiotics with different activities.

### Integrated genome mining leads to identification of mycofactocin-like BGCs in *Saccharopolyspora*

On top of known resistance gene models, ARTS2 reference set includes phylum-specific essential housekeeping genes (core models). Both models are checked for duplication, horizontal gene transfer, and localization within a BGC, which signifies potential bioactivity (25). In this section, we carried out additional exploratory analysis for a selected subnetwork of 17 BGCs of ranthipeptide type with unknown functions. These 17 BGCs do not have a match through KnownClusterBlast and BiGFAM-DB, but all shared a hit against an ARTS2 core model TIGR03997 (Figure 7A). The TIGR03997 model was identified as a mycofactocin system oxidoreductase, a related biosynthetic component of a known redox cofactor in *Mycobacterium tuberculosis* (88). Even though this profile does not satisfy several ARTS criteria for self-resistance, it indicated that the detected BGCs might code for mycofactocin-like redox cofactors.

**Figure 7.**
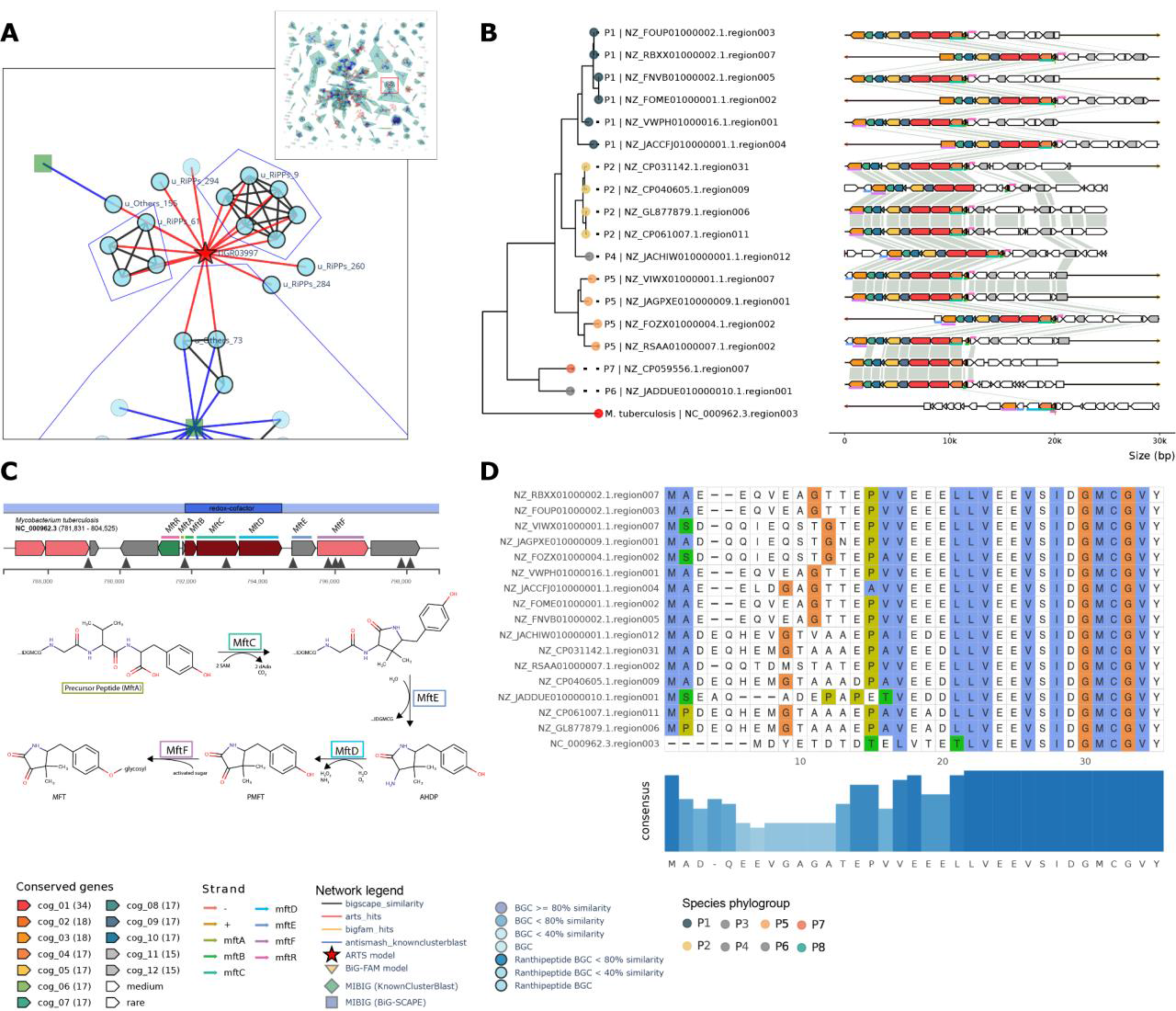
Identification of mycofactocin-like BGCs in *Saccharopolyspora* through integrated genome mining with ARTS2 A) Integration of antiSMASH KnownClusterBlast, ARTS2, BiG-FAM hits, and BiG-SCAPE sequence similarity networks enables quick exploration and visualization for unknown BGCs with potential bioactivities. The zoomed panel shows a group of Ranthipeptide BGCs with low KnownClusterBlast similarity score with hits to ARTS2 model (TIGR03997). B) Cluster comparison of the putative Mycofactocin BGCs (Data S7), classified as redox-cofactor type in antiSMASH, as described in (88). Alignment shows conserved RS-SPASM (MftC) and Mycofactocin RRE (MftB) signature, but missing TIGR03966 profile in *Saccharopolyspora*. C) the proposed Mycofactocin biosynthesis pathway in *Mycobacterium tuberculosis,* adapted from (91) D) Mycofactocin precursor peptide (MftA) comparison in *Saccharopolyspora* showed distinct conserved C-terminal residues (–IDGMCGVY).

We did a further investigation into these 17 candidate BGCs by creating a new PEP with BGC snakefile (Data S7) and comparing the detected BGCs with a mycofactocin BGC from *Mycobacterium tuberculosis*. We found that the majority of the genes showed sequence similarities to at least six out of the seven genes involved in mycofactocin biosynthesis (Figure 7B and C), with the exception of *mftD*. Multiple sequence alignment of the candidate precursor peptides (*mftA*) exhibited the characteristic C-terminal pattern (IDGMCGVY) associated with mycofactocin (Figure 7D) (88). The missing gene signature (*mftD*) encodes an oxidoreductase (TIGR03966) that is required for the synthesis of pre-mycofactocin through oxidative deamination of *3-amino-5-[(p-hydroxyphenyl)methyl]-4,4-dimethyl-2-pyrrolidinone* (AHDP) (88). The *Saccharapolyspora* BGCs contained a gene without signficant similarity to mftD but a hit against the ARTS profile TIGR03997, which is annotated as FadH/OYE family oxidoreductase with unknown function. Nevertheless, experimental evidence showed that *S. erythraea* is able to produce a methylated variant of Mycofactocin (89). These findings strongly suggest that the investigated BGCs may be responsible for producing a compound that closely resembles mycofactocin, instead of a ranthipeptide as was initially discovered with the rule-based logic of antiSMASH. The classification discrepancy was likely due to the absence of the TIGR03966 (MftD) profile in the candidate BGCs, which are required by the antiSMASH version 6.1.1 rule to recognize redox cofactor BGCs. The missing profile then registers the BGC region as ranthipeptide, even though the clusters don’t seem to indicate the six Cys in forty-five residues (SCIFF) pattern in the precursor peptides (90). Hence, the BGCFlow framework provides a global overview and guidelines for investigating BGCs and GCFs in finer detail that can reveal more knowledge about the BGCs of interest.

## DISCUSSION

Genome mining tools like antiSMASH have revealed the significantly higher biosynthetic potential of various bacteria to produce novel secondary metabolites with various applications in medicine and industrial biotechnology (15). In recent years, the search for novel natural products with bioactivity is evolving from purely chemical and activity screening-based approaches to genome analysis-assisted discovery (14, 92). This paradigm shift in the era of pangenome availability will require scientists to deploy comprehensive platforms to organize the large datasets, metadata, and results in a findable, accessible, interoperable, and reusable (FAIR) way. The data workflow management systems and databases provide effective solutions to organize large genome mining projects. Here, the BGCFlow demonstrates the first end-to-end analysis platform to unify large-scale BGC and genome data, increasing the support for FAIR data principles.

Currently, workflows such as AQUAMIS, ASA^3^P, TORMES, Bactopia, and others automate many of the “generic” assembly, annotation, and pangenome analysis software (29, 36–38), but provide no information on specialized metabolite BGCs. Many of the genome mining tools like antiSMASH, BiG-SCAPE, and BiG-SLICE allow the detection of BGCs and GCFs, whereas databases like antiSMASH-DB, BiG-FAM, MIBIG provide precalculated BGCs, GCFs, or catalog of known BGCs (21, 26–28, 69, 93). With a primary focus on integrating secondary/specialized metabolite genome mining resources under one platform, BGCFlow assists in automating the computational analysis in scalable, adaptable, reiterable fashion for various small to large-scale customized projects.

Particularly, any exploratory data analysis requires scientists to dynamically change the dataset, adding or removing samples and running different analytical tools as the investigation continues. The notebooks and the database provided here would be a starting point for many users to expand the exploratory analysis for the design of the study. Inspired by the cookiecutter data science project (https://drivendata.github.io/cookiecutter-data-science), BGCFlow data structure, combined with Snakemake, enables efficient re-use of data via input re-entries. While the notebooks are meant for flexible exploratory analysis, it is recommended to utilize the combined PEP configuration and Snakemake for reproducibility. On top of the main workflow, users can also utilize sub-workflows for direct comparison of BGCs (as demonstrated with examples of erythreapeptin and mycofactocin comparison) or write a custom pipeline.

Comparing BGCs and grouping them into functionally related GCFs remains a key challenge in genome mining studies (26, 27, 94–97). The accuracy of such comparisons can be influenced by several factors, depending on the scale and quality of the dataset. i) Relying solely on gene sequence similarity for comparisons can lead to the grouping together of distinct BGCs that share repetitive sequences. ii) Comparing hybrid regions of BGCs, which may contain multiple independent BGCs, can impact the assessment of similarity. iii) The precise determination of BGC boundaries for comparison purposes is challenging. iv) Poor assembly quality frequently results in BGCs being fragmented. v) Effectively navigating and identifying relevant GCFs from a vast number of clusters demands the implementation of efficient prioritization strategies, as not all tools are equipped to handle larger datasets. By leveraging the knowledge extracted from antiSMASH KnownClusterBlast, MIBIG-DB, BiG-SCAPE, BiG-FAM DB, and ARTS into an enriched sequence similarity network, BGCFlow enables the more accurate definition of GCFs and concise comparative analysis of BGCs.

As a case study, we ran the BGCFlow on a set of publicly available genomes of *Saccharopolyspora* genus, which is known to produce several industrially relevant natural products with still underexplored bioactive potential (73). Various steps of the analysis carried out on different subsets highlight the reiterable nature of BGCFlow analysis. The phylogenetic analysis provides a guideline for efficiently investigating the distribution of different BGCs across phylogroups, for example different classes of lanthipeptides. The calculated enriched similarity network of the BGCs with different genome mining resources improved classification of BGCs and GCFs compared to the individual tools. Our analysis revealed more accurate identification of spinosyn-like, erythreapeptin-like, and mycofactin-like GCFs through connections to KnownClusterBlast based MIBiG BGCs or ARTS profiles. For instance, when classifying spinosyn-like BGCs using BiG-SCAPE with a cutoff of 0.3, we encountered challenges due to multiple BGCs residing in the same genomic region, resulting in their classification into different GCFs. In another example, the integration of knowledge from ARTS allowed us to re-evaluate the antiSMASH rule-based classification of ranthipeptides. Our analysis uncovered 17 ranthipeptide BGCs actually resembling mycofactocin biosynthesis, a redox-cofactor type also recently identified to be produced by *Saccharapolyspora* (89). These findings highlight the importance of not only integration but also the need to carefully evaluate accuracy of each predicted GCF as per the case. While literature curated databases like MIBIG are valuable resources, they may still contain incomplete entries and significant gaps to current knowledge, especially considering the ever-growing volume of publications in the natural product field.

Despite several advances in genome mining tools over the last decade, it is important to note that many of the BGC related analyses and predictions still require experimental validations. Therefore, it is crucial to exercise caution and critically interpret the data-driven predictions. Nevertheless, analyzing large genomic datasets, even with domain knowledge expertise, presents challenges due to variations in data quality, software versions, and database inconsistencies. To address these issues, BGCFlow aims to provide a scalable, explorable and reproducible platform for integrated genome mining analysis. By following workflow standards and incorporating data engineering best practices, BGCFlow provides a dynamic environment that facilitates not only the execution of various analysis tools but also enables critical exploratory analysis of BGCs at different scales of pangenome-level, BGC-level and sequence-level. Thus, we expect that the BGCFlow will streamline the research process along with facilitating collaboration and knowledge sharing among researchers working in the genome mining field.

## MATERIALS AND METHODS

### Installing and running BGCFlow using the BGCFlow wrapper

BGCFlow software environment consists of three components: (1) a snakemake workflow (https://github.com/NBChub/bgcflow), (2) a DBT pipeline for extraction of tabular data (https://github.com/NBChub/bgcflow_dbt-duckdb), and (3) a python command line interface called *bgcflow_wrapper* (https://github.com/NBChub/bgcflow_wrapper). The wrapper can be installed using PIP directly from the repository (*pip install git+*https://github.com/NBChub/bgcflow_wrapper.git), and has the command lines to access all of the features in BGCFlow.

### Design of the Snakemake rules for external software

As part of the Snakemake workflow, several external computational tools were selected for different stages of the analysis. In the first step, the installation guide from the individual software’s repository was followed to create a conda-based environment using *yaml* files stored in the *workflow/envs/* directory. The version of all dependencies and the main software can be changed in these environment files. For a few cases, additional rules were created for the installation, which typically involved downloading additional resources and databases. These additional resources are stored in the *resources* directory, and users can also provide the pre-downloaded resource directories in the main configuration yaml file. A typical script for the Snakemake rule includes paths to the conda environment, inputs and outputs of the software in the interim directory, along with the required parameters of the command line.

### Customized Snakemake rules data wrangling

To seamlessly run various software included in BGCFlow, many customized rules were generated that make sure the formatting of input and output files were interoperable (Table S2). Such data wrangling typically included rules that prepare the input formatting and process the outputs in tabular interoperable format stored in the *processed* directory of each project. These data-wrangling rules involved running custom Python scripts from the bgcflow Python package directory. Many of the customized rules also involved additional downstream analysis of visualizing data.

### Visualizing Jupyter Notebook reports

Each of the major rules is supported with a templated Jupyter Notebook for exploratory analysis of the results. These templated notebooks can be run with the *bgcflow serve* command, which copies the template notebook from the *workflow/notebooks* directory to the *processed* directory of each project. These template notebooks are meant to be a starting point for exploratory analysis and thus can be edited and expanded for each project. The notebooks are then converted to mkdocs reports corresponding to each rule. The online server provides navigation of the interactive reports for each rule. The reports are also connected to the main results and tables, providing options for downloads for downstream exploration. For example, the rules for antiSMASH, BiG-SCAPE, and BiG-SLICE provide links to HTML reports provided by these individual tools.

### Exporting results tables in OLAP database

Once the BGCFlow run is complete, all tables from various pipelines are stored as Apache Parquet files for efficient data storage. The command *bgcflow build* enables the ability to extract, load, and transform the data into SQL databases through the use of Data Build Tools (DBT). Here, a set of SQL schemas are generated to easily interconnect BGC and genome information with other analysis and metadata for data queries (Figure S4). By default, BGCFlow utilizes DuckDB, an embedded Online Analytical Processing (OLAP) database management system. Users can create additional data models and use other database management systems by modifying DBT profiles (https://github.com/NBChub/bgcflow_dbt-duckdb). For an interactive exploration of the database, the command *bgcflow serve –metabase* will start up a Metabase server, a popular business intelligence and visualization tool.

### Genome mining and comparative analysis of *Saccharopolyspora* BGCs

This study incorporates several BGCFlow projects, defined in the PEP formats (available at Zenodo: https://doi.org/10.5281/zenodo.8018055) as follows:

#### PEP definition for all quality Saccharopolospora genomes

We searched for the *Saccharopolyspora* genus at NCBI RefSeq with filters to include all assembly levels of the complete genome, chromosome, scaffold, and contig on 25 October 2022, resulting in the list of 42 accession IDs. The sample table with these 42 accession IDs was provided in the PEP configuration initiated with the project name *qc_saccharopolyspora*, which aimed to carry out the quality check and taxonomic placement (Data S1). The selected BGCFlow rules for this first example PEP included: *checkm*, *gtdbtk*, *seqfu*, and *mash*. The results of the run were investigated using the Jupyter notebooks for each rule provided in the directory *data/processed/qc_saccharopolyspora/docs/* and visualized with reports using *bgcflow serve -- project qc_saccharopolyspora* command. Based on the assembly quality, the genomes were categorized into three groups: 1) HQ (high-quality genomes of complete genome or chromosome level assembly as defined by NCBI RefSeq), 2) MQ (medium-quality genomes with less than 50 contigs, N50 score higher than 100 Kbps), and 3) LQ (remaining low-quality genomes) (Figure S6, Figure 3a). The generated GTDB taxonomy definitions were used for the downstream analysis. The MASH-distance calculation was used to group 42 genomes in 7 different clusters of genomes called phylogroups in this study.

#### PEP definition for medium to high-quality Saccharopolospora genomes

To demonstrate full-scale BGCFlow application, we selected 26 genomes from HQ and MQ categories as defined above in the second PEP configuration initiated with the project name *mq_saccharopolyspora* (Data S2). All 19 rules were selected for the BGCFlow run of this PEP. The results of each rule are analyzed using the custom Jupyter notebooks stored in the directory *data/processed/mq_saccharopolyspora/docs*. The autoMLST-based phylogenetic tree of all 26 genomes was used for downstream investigation including phylogenetic distribution of BGCs using the *ggtree* R package (67). The MASH-based clustering was used to group the genomes into 7 phylogroups that were used across various visualizations. The BGC comparative analysis was carried out by BiG-SLICE and BiG-SCAPE and the networks were further visualized using Plotly (Figure 4).

### Enriched GCF Network Exploration

The enriched similarity network of BGCs were built on top of the resulting BiG-SCAPE sequence similarity network by adding nodes and edges from the result of ARTS, KnownClusterBlast, and query against BiG-FAM database using the python package networkx. MIBIG entries from KnownClusterBlast, ARTS model, and BiG-FAM models were added as nodes, while new edges were created to connect BGC nodes with the newly introduced nodes. Shannon Index (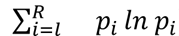) for each BiG-FAM models were calculated at the genus level, where *p_i_* is the proportion of the *i*-th genus in the model. BiG-FAM models 202087, 200946, 210179, 213140, 201682, 201608, 205957, 215277, 201830, and 202082 were removed from the network based on the low percentage of the assigned top genus (<30%) and high Shannon Index. Clusters of connected components were calculated using networkx but ignoring ARTS nodes. For clusters with *n* > 2, a convex hull was drawn using the package scipy spatial.

### Comparison of Gene Cluster Families

Sets of selected GCFs and BGCs were defined in a PEP format and were run using a comparative BGC analysis sub-workflow (*bgcflow run –snakefile workflow/BGC*) and further refined in Jupyter notebook scripts (https://github.com/NBChub/saccharopolyspora_manuscrip). The sub-workflow included specific tools for an in-depth comparison of the BGC structure of sequence alignments. Briefly, MMSeqs2 was used to build a protein sequence similarity network of selected BGCs, which allowed us to cluster the coding sequences into candidate Clusters of Orthologous Groups of proteins (COGs) (98). A phylogenetic tree was built by concatenated alignment of single-copy orthologs which are conserved across all selected BGCs using Seqkit (99) and IQTree (100). The resulting phylogenetic tree and sequence similarity network are then visualized using the R package GGGenomes (https://github.com/thackl/gggenomes) (101) and GGTree (67). The gene comparison was enriched with CBlaster hits (65) to a selected reference BGCs and links between genes are made using CLinker (102). Multiple sequence alignments of the candidate precursor peptides are built using Clustal Omega (103) and visualized using the GGMSA R package (104).

## AVAILABILITY

BGCFlow software environment consists of three components: a snakemake workflow (https://github.com/NBChub/bgcflow), a DBT pipeline for extraction of tabular data (https://github.com/NBChub/bgcflow_dbt-duckdb), and a python command line interface called bgcflow_wrapper (https://github.com/NBChub/bgcflow_wrapper). The Jupyter Notebooks for the data analysis are available on GitHub at https://github.com/NBChub/saccharopolyspora_manuscript. The data of the entire BGCFlow run on the *Saccharopolyspora* genus is available at Zenodo: https://doi.org/10.5281/zenodo.8018055.

## SUPPLEMENTARY DATA

Supplementary Data are available at Zenodo: https://doi.org/10.5281/zenodo.8018055.

## Supporting information

ZIP archive of supplementary data tables (MS Excel)

## ACKNOWLEDGEMENT

We thank Kai Blin, Simon Shaw, Tue S. Jorgensen, Emre Özdemir, Roberto S. Navarro, Matiss Maleckis, Lasse J. D. Nielsen, Pernille K. Bech, Eva B. Sterndorff, and Zhijie Yang for important discussion and feedback during the development and testing of BGCFlow.

## References

1. Fullam,A., Letunic,I., Schmidt,T.S.B., Ducarmon,Q.R., Karcher,N., Khedkar,S., Kuhn,M., Larralde,M., Maistrenko,O.M., Malfertheiner,L., et al. (2022) proGenomes3: approaching one million accurately and consistently annotated high-quality prokaryotic genomes. Nucleic Acids Res., 10.1093/nar/gkac1078.

2. Doron,S., Melamed,S., Ofir,G., Leavitt,A., Lopatina,A., Keren,M., Amitai,G. and Sorek,R. (2018) Systematic discovery of antiphage defense systems in the microbial pangenome. Science, 359, 1–11.

3. Zhang,Y., Zhang,H., Zhang,Z., Qian,Q., Zhang,Z. and Xiao,J. (2022) ProPan: a comprehensive database for profiling prokaryotic pan-genome dynamics. Nucleic Acids Res., 10.1093/nar/gkac832.

4. Hyun,J.C., Monk,J.M. and Palsson,B.O. (2022) Comparative pangenomics: analysis of 12 microbial pathogen pangenomes reveals conserved global structures of genetic and functional diversity. BMC Genomics, 23, 7.

5. Abram,K., Udaondo,Z., Bleker,C., Wanchai,V., Wassenaar,T.M., Robeson,M.S.,2nd and Ussery,D.W. (2021) Mash-based analyses of Escherichia coli genomes reveal 14 distinct phylogroups. Commun Biol, 4, 117.

6. Mageiros,L., Méric,G., Bayliss,S.C., Pensar,J., Pascoe,B., Mourkas,E., Calland,J.K., Yahara,K., Murray,S., Wilkinson,T.S., et al. (2021) Genome evolution and the emergence of pathogenicity in avian Escherichia coli. Nat. Commun., 12, 765.

7. Mohite,O.S., Lloyd,C.J., Monk,J.M., Weber,T. and Palsson,B.O. (2022) Pangenome analysis of Enterobacteria reveals richness of secondary metabolite gene clusters and their associated gene sets. Synth Syst Biotechnol, 7, 900–910.

8. Shi,Y.-M., Hirschmann,M., Shi,Y.-N., Ahmed,S., Abebew,D., Tobias,N.J., Grün,P., Crames,J.J., Pöschel,L., Kuttenlochner,W., et al. (2022) Global analysis of biosynthetic gene clusters reveals conserved and unique natural products in entomopathogenic nematode-symbiotic bacteria. Nat. Chem., 14, 701–712.

9. Kloosterman,A.M., Cimermancic,P., Elsayed,S.S., Du,C., Hadjithomas,M., Donia,M.S., Fischbach,M.A., van Wezel,G.P. and Medema,M.H. (2020) Expansion of RiPP biosynthetic space through integration of pan-genomics and machine learning uncovers a novel class of lanthipeptides. PLOS Biology, 18, e3001026.

10. Wright,G.D. (2017) Opportunities for natural products in 21st century antibiotic discovery. Nat. Prod. Rep., 34, 694–701.

11. Huang,M., Lu,J.-J. and Ding,J. (2021) Natural Products in Cancer Therapy: Past, Present and Future. Nat. Products Bioprospect., 11, 5–13.

12. Atanasov,A.G., Zotchev,S.B., Dirsch,V.M., International Natural Product Sciences Taskforce and Supuran, C.T. (2021) Natural products in drug discovery: advances and opportunities. Nat. Rev. Drug Discov., 20, 200–216.

13. Medema,M.H. (2021) The year 2020 in natural product bioinformatics: an overview of the latest tools and databases. Nat. Prod. Rep., 38, 301–306.

14. Ziemert,N., Alanjary,M. and Weber,T. (2016) The evolution of genome mining in microbes - a review. Nat. Prod. Rep., 33, 988–1005.

15. Gavriilidou,A., Kautsar,S.A., Zaburannyi,N., Krug,D., Müller,R., Medema,M.H. and Ziemert,N. (2022) Compendium of specialized metabolite biosynthetic diversity encoded in bacterial genomes. Nat Microbiol, 7, 726–735.

16. Steinke,K., Mohite,O.S., Weber,T. and Kovács,Á.T. (2021) Phylogenetic Distribution of Secondary Metabolites in the Bacillus subtilis Species Complex. mSystems, 6.

17. Adamek,M., Alanjary,M., Sales-Ortells,H., Goodfellow,M., Bull,A.T., Winkler,A., Wibberg,D., Kalinowski,J. and Ziemert,N. (2018) Comparative genomics reveals phylogenetic distribution patterns of secondary metabolites in Amycolatopsis species. BMC Genomics, 19, 426.

18. Chase,A.B., Sweeney,D., Muskat,M.N., Guillén-Matus,D.G. and Jensen,P.R. (2021) Vertical Inheritance Facilitates Interspecies Diversification in Biosynthetic Gene Clusters and Specialized Metabolites. MBio, 12, e0270021.

19. Medema,M.H., Cimermancic,P., Sali,A., Takano,E. and Fischbach,M.A. (2014) A systematic computational analysis of biosynthetic gene cluster evolution: lessons for engineering biosynthesis. PLoS Comput. Biol., 10, e1004016.

20. Donia,M.S., Cimermancic,P., Schulze,C.J., Wieland Brown,L.C., Martin,J., Mitreva,M., Clardy,J., Linington,R.G. and Fischbach,M.A. (2014) A systematic analysis of biosynthetic gene clusters in the human microbiome reveals a common family of antibiotics. Cell, 158, 1402–1414.

21. Blin,K., Shaw,S., Kloosterman,A.M., Charlop-Powers,Z., van Wezel,G.P., Medema,M.H. and Weber,T. (2021) antiSMASH 6.0: improving cluster detection and comparison capabilities. Nucleic Acids Res., 49, W29–W35.

22. Blin,K., Shaw,S., Augustijn,H.E., Reitz,Z.L., Biermann,F., Alanjary,M., Fetter,A., Terlouw,B.R., Metcalf,W.W., Helfrich,E.J.N., et al. (2023) antiSMASH 7.0: new and improved predictions for detection, regulation, chemical structures and visualisation. Nucleic Acids Res., 10.1093/nar/gkad344.

23. Blin,K., Medema,M.H., Kottmann,R., Lee,S.Y. and Weber,T. (2017) The antiSMASH database, a comprehensive database of microbial secondary metabolite biosynthetic gene clusters. Nucleic Acids Res., 45, D555–D559.

24. Kautsar,S.A., Blin,K., Shaw,S., Navarro-Muñoz,J.C., Terlouw,B.R., van der Hooft,J.J.J., van Santen,J.A., Tracanna,V., Suarez Duran,H.G., Pascal Andreu,V., et al. (2020) MIBiG 2.0: a repository for biosynthetic gene clusters of known function. Nucleic Acids Res., 48, D454–D458.

25. Mungan,M.D., Alanjary,M., Blin,K., Weber,T., Medema,M.H. and Ziemert,N. (2020) ARTS 2.0: feature updates and expansion of the Antibiotic Resistant Target Seeker for comparative genome mining. Nucleic Acids Res., 48, W546–W552.

26. Navarro-Muñoz,J.C., Selem-Mojica,N., Mullowney,M.W., Kautsar,S.A., Tryon,J.H., Parkinson,E.I., De Los Santos,E.L.C., Yeong,M., Cruz-Morales,P., Abubucker,S., et al. (2020) A computational framework to explore large-scale biosynthetic diversity. Nat. Chem. Biol., 16, 60–68.

27. Kautsar,S.A., van der Hooft,J.J.J., de Ridder,D. and Medema,M.H. (2021) BiG-SLiCE: A highly scalable tool maps the diversity of 1.2 million biosynthetic gene clusters. Gigascience, 10.

28. Kautsar,S.A., Blin,K., Shaw,S., Weber,T. and Medema,M.H. (2021) BiG-FAM: the biosynthetic gene cluster families database. Nucleic Acids Res., 49, D490–D497.

29. Petit,R.A.,3rd and Read,T.D. (2020) Bactopia: a Flexible Pipeline for Complete Analysis of Bacterial Genomes. mSystems, 5.

30. Cornwell,M., Vangala,M., Taing,L., Herbert,Z., Köster,J., Li,B., Sun,H., Li,T., Zhang,J., Qiu,X., et al. (2018) VIPER: Visualization Pipeline for RNA-seq, a Snakemake workflow for efficient and complete RNA-seq analysis. BMC Bioinformatics, 19, 135.

31. Köster,J. and Rahmann,S. (2012) Snakemake—a scalable bioinformatics workflow engine. Bioinformatics, 28, 2520–2522.

32. Di Tommaso,P., Chatzou,M., Floden,E.W., Barja,P.P., Palumbo,E. and Notredame,C. (2017) Nextflow enables reproducible computational workflows. Nat. Biotechnol., 35, 316–319.

33. Voss,K., Van der Auwera,G. and Gentry,J. Full-stack genomics pipelining with GATK4+ WDL+ Cromwell. F1000Res.

34. Wratten,L., Wilm,A. and Göke,J. (2021) Reproducible, scalable, and shareable analysis pipelines with bioinformatics workflow managers. Nat. Methods, 18, 1161–1168.

35. Chevrette,M.G. and Handelsman,J. (2021) Needles in haystacks: reevaluating old paradigms for the discovery of bacterial secondary metabolites. Nat. Prod. Rep., 38, 2083–2099.

36. Deneke,C., Brendebach,H., Uelze,L., Borowiak,M., Malorny,B. and Tausch,S.H. (2021) Species-Specific Quality Control, Assembly and Contamination Detection in Microbial Isolate Sequences with AQUAMIS. Genes, 12.

37. Schwengers,O., Hoek,A., Fritzenwanker,M., Falgenhauer,L., Hain,T., Chakraborty,T. and Goesmann,A. (2020) ASA3P: An automatic and scalable pipeline for the assembly, annotation and higher-level analysis of closely related bacterial isolates. PLoS Comput. Biol., 16, e1007134.

38. Quijada,N.M., Rodríguez-Lázaro,D., Eiros,J.M. and Hernández,M. (2019) TORMES: an automated pipeline for whole bacterial genome analysis. Bioinformatics, 35, 4207–4212.

39. Sayers,E.W., Bolton,E.E., Brister,J.R., Canese,K., Chan,J., Comeau,D.C., Farrell,C.M., Feldgarden,M., Fine,A.M., Funk,K., et al. (2023) Database resources of the National Center for Biotechnology Information in 2023. Nucleic Acids Res., 51, D29–D38.

40. Wattam,A.R., Abraham,D., Dalay,O., Disz,T.L., Driscoll,T., Gabbard,J.L., Gillespie,J.J., Gough,R., Hix,D., Kenyon,R., et al. (2014) PATRIC, the bacterial bioinformatics database and analysis resource. Nucleic Acids Res., 42, D581–91.

41. Sheffield,N.C., Stolarczyk,M., Reuter,V.P. and Rendeiro,A.F. (2021) Linking big biomedical datasets to modular analysis with Portable Encapsulated Projects. Gigascience, 10.

42. Chaumeil,P.-A., Mussig,A.J., Hugenholtz,P. and Parks,D.H. (2019) GTDB-Tk: a toolkit to classify genomes with the Genome Taxonomy Database. Bioinformatics, 36, 1925– 1927.

43. Seemann,T. (2014) Prokka: rapid prokaryotic genome annotation. Bioinformatics, 30, 2068– 2069.

44. Panoptes: Monitor computational workflows in real time https://github.com/panoptes-organization/panoptes.

45. Raasveldt,M. and Mühleisen,H. (2019) DuckDB: an Embeddable Analytical Database. In Proceedings of the 2019 International Conference on Management of Data, SIGMOD’19. Association for Computing Machinery, New York, NY, USA, pp. 1981–1984.

46. Yang,J., Liu,Y., Shang,J., Huang,Y., Yu,Y., Li,Z., Shi,L. and Ran,Z. (2022) BioVisReport: A Markdown-based lightweight website builder for reproducible and interactive visualization of results from peer-reviewed publications. Comput. Struct. Biotechnol. J., 20, 3133–3139.

47. Vink,T. (2022) Reproducible Reports with MkDocs. https://www.timvink.nl//reproducible-reports-with-mkdocs/.

48. Telatin,A., Fariselli,P. and Birolo,G. (2021) SeqFu: A Suite of Utilities for the Robust and Reproducible Manipulation of Sequence Files. Bioengineering (Basel), 8.

49. Sánchez-Navarro,R., Nuhamunada,M., Mohite,O.S., Wasmund,K., Albertsen,M., Gram,L., Nielsen,P.H., Weber,T. and Singleton,C.M. (2022) Long-Read Metagenome-Assembled Genomes Improve Identification of Novel Complete Biosynthetic Gene Clusters in a Complex Microbial Activated Sludge Ecosystem. mSystems, 7, e0063222.

50. Parks,D.H., Imelfort,M., Skennerton,C.T., Hugenholtz,P. and Tyson,G.W. (2015) CheckM: assessing the quality of microbial genomes recovered from isolates, single cells, and metagenomes. Genome Res., 25, 1043–1055.

51. Bowers,R.M., Kyrpides,N.C., Stepanauskas,R., Harmon-Smith,M., Doud,D., Reddy,T.B.K., Schulz,F., Jarett,J., Rivers,A.R., Eloe-Fadrosh,E.A., et al. (2017) Minimum information about a single amplified genome (MISAG) and a metagenome-assembled genome (MIMAG) of bacteria and archaea. Nat. Biotechnol., 35, 725–731.

52. Parks,D.H., Chuvochina,M., Rinke,C., Mussig,A.J., Chaumeil,P.-A. and Hugenholtz,P. (2022) GTDB: an ongoing census of bacterial and archaeal diversity through a phylogenetically consistent, rank normalized and complete genome-based taxonomy. Nucleic Acids Res., 50, D785–D794.

53. Ondov,B.D., Treangen,T.J., Melsted,P., Mallonee,A.B., Bergman,N.H., Koren,S. and Phillippy,A.M. (2016) Mash: fast genome and metagenome distance estimation using MinHash. Genome Biol., 17, 132.

54. Jain,C., Rodriguez-R,L.M., Phillippy,A.M., Konstantinidis,K.T. and Aluru,S. (2018) High throughput ANI analysis of 90K prokaryotic genomes reveals clear species boundaries. Nat. Commun., 9, 5114.

55. Hyatt,D., Chen,G.-L., Locascio,P.F., Land,M.L., Larimer,F.W. and Hauser,L.J. (2010) Prodigal: prokaryotic gene recognition and translation initiation site identification. BMC Bioinformatics, 11, 119.

56. Cantalapiedra,C.P., Hernández-Plaza,A., Letunic,I., Bork,P. and Huerta-Cepas,J. (2021) eggNOG-mapper v2: Functional Annotation, Orthology Assignments, and Domain Prediction at the Metagenomic Scale. Mol. Biol. Evol., 38, 5825–5829.

57. Devoid,S., Overbeek,R., DeJongh,M., Vonstein,V., Best,A.A. and Henry,C. (2013) Automated genome annotation and metabolic model reconstruction in the SEED and Model SEED. Methods Mol. Biol., 985, 17–45.

58. Hernández-Plaza,A., Szklarczyk,D., Botas,J., Cantalapiedra,C.P., Giner-Lamia,J., Mende,D.R., Kirsch,R., Rattei,T., Letunic,I., Jensen,L.J., et al. (2023) eggNOG 6.0: enabling comparative genomics across 12 535 organisms. Nucleic Acids Res., 51, D389–D394.

59. Galperin,M.Y., Wolf,Y.I., Makarova,K.S., Vera Alvarez,R., Landsman,D. and Koonin,E.V. (2021) COG database update: focus on microbial diversity, model organisms, and widespread pathogens. Nucleic Acids Res., 49, D274–D281.

60. The Gene Ontology resource: enriching a GOld mine (2021) Nucleic Acids Res., 49, D325–D334.

61. Kanehisa,M., Furumichi,M., Sato,Y., Kawashima,M. and Ishiguro-Watanabe,M. (2023) KEGG for taxonomy-based analysis of pathways and genomes. Nucleic Acids Res., 51, D587–D592.

62. Page,A.J., Cummins,C.A., Hunt,M., Wong,V.K., Reuter,S., Holden,M.T.G., Fookes,M., Falush,D., Keane,J.A. and Parkhill,J. (2015) Roary: rapid large-scale prokaryote pan genome analysis. Bioinformatics, 31, 3691–3693.

63. Kim,G.B., Gao,Y., Palsson,B.O. and Lee,S.Y. (2021) DeepTFactor: A deep learning-based tool for the prediction of transcription factors. Proc. Natl. Acad. Sci. U. S. A., 118.

64. Buchfink,B., Reuter,K. and Drost,H.-G. (2021) Sensitive protein alignments at tree-of-life scale using DIAMOND. Nat. Methods, 18, 366–368.

65. Gilchrist,C.L.M., Booth,T.J., van Wersch,B., van Grieken,L., H Medema,M. and Chooi,Y.-H. (2021) cblaster: a remote search tool for rapid identification and visualization of homologous gene clusters. Bioinformatics Advances, 1, 1–10.

66. Alanjary,M., Steinke,K. and Ziemert,N. (2019) AutoMLST: an automated web server for generating multi-locus species trees highlighting natural product potential. Nucleic Acids Res., 47, W276–W282.

67. Yu,G., Smith,D.K., Zhu,H., Guan,Y. and Lam,T.T.-Y. (2017) Ggtree : An r package for visualization and annotation of phylogenetic trees with their covariates and other associated data. Methods Ecol. Evol., 8, 28–36.

68. Letunic,I. and Bork,P. (2019) Interactive Tree Of Life (iTOL) v4: recent updates and new developments. Nucleic Acids Res., 47, W256–W259.

69. Terlouw,B.R., Blin,K., Navarro-Muñoz,J.C., Avalon,N.E., Chevrette,M.G., Egbert,S., Lee,S., Meijer,D., Recchia,M.J.J., Reitz,Z.L., et al. (2022) MIBiG 3.0: a community-driven effort to annotate experimentally validated biosynthetic gene clusters. Nucleic Acids Res., 51, gkac1049.

70. Caicedo-Montoya,C., Manzo-Ruiz,M. and Ríos-Estepa,R. (2021) Pan-Genome of the Genus Streptomyces and Prioritization of Biosynthetic Gene Clusters With Potential to Produce Antibiotic Compounds. Frontiers in Microbiology, 12.

71. Otani,H., Udwary,D.W. and Mouncey,N.J. (2022) Comparative and pangenomic analysis of the genus Streptomyces. Sci. Rep., 12, 18909.

72. Letzel,A.-C., Li,J., Amos,G.C.A., Millán-Aguiñaga,N., Ginigini,J., Abdelmohsen,U.R., Gaudêncio,S.P., Ziemert,N., Moore,B.S. and Jensen,P.R. (2017) Genomic insights into specialized metabolism in the marine actinomycete Salinispora. Environ. Microbiol., 19, 3660–3673.

73. Sayed,A.M., Abdel-Wahab,N.M., Hassan,H.M. and Abdelmohsen,U.R. (2020) Saccharopolyspora: an underexplored source for bioactive natural products. J. Appl. Microbiol., 128, 314–329.

74. Ma,D., Liu,S., Liu,H., Nan,M., Xu,Y., Han,X. and Mao,J. (2022) Developing an innovative raw wheat Qu inoculated with Saccharopolyspora and its application in Huangjiu. J. Sci. Food Agric., 102, 7301–7312.

75. Garrod,L.P. (1957) The erythromycin group of antibiotics. Br. Med. J., 2, 57–63.

76. Kirst,H.A., Michel,K.H., Martin,J.W., Creemer,L.C., Chio,E.H., Yao,R.C., Nakatsukasa,W.M., Boeck,L.D., Occolowitz,J.L., Paschal,J.W., et al. (1991) A83543A-D, unique fermentation-derived tetracyclic macrolides. Tetrahedron Lett., 32, 4839–4842.

77. Sparks,T.C., Crouse,G.D. and Durst,G. (2001) Natural products as insecticides: the biology, biochemistry and quantitative structure–activity relationships of spinosyns and spinosoids. Pest Manag. Sci., 57, 896–905.

78. Saygin,H., Ay,H., Guven,K., Inan-Bektas,K., Cetin,D. and Sahin,N. (2021) Saccharopolyspora karakumensis sp. nov., Saccharopolyspora elongata sp. nov., Saccharopolyspora aridisoli sp. nov., Saccharopolyspora terrae sp. nov. and their biotechnological potential revealed by genome analysis. Syst. Appl. Microbiol., 44, 126270.

79. Lacey,J., Goodfellow,M., Lacy,J. and Goodfellow,M. (1975) A novel actinomycete from sugar-cane bagasse: Saccharopolyspora hirsuta gen. et. sp. nov. J. Gen. Microbiol., 88, 75–85.

80. Reimer,L.C., Sardà Carbasse,J., Koblitz,J., Ebeling,C., Podstawka,A. and Overmann,J. (2022) BacDive in 2022: the knowledge base for standardized bacterial and archaeal data. Nucleic Acids Res., 50, D741–D746.

81. Goodfellow,M., Lacey,J., Athalye,M., Embley,T.M. and Bowen,T. (1989) Saccharopolyspora gregorii and Saccharopolyspora hordei: Two New Actinomycete Species from Fodder. Microbiology, 135, 2125–2139.

82. Intra,B., Euanorasetr,J., Také,A., Inahashi,Y., Mori,M., Panbangred,W. and Matsumoto,A. (2019) Saccharopolyspora rhizosphaerae sp. nov., an actinomycete isolated from rhizosphere soil in Thailand. Int. J. Syst. Evol. Microbiol., 69, 1299–1305.

83. Dale,S.E., Doherty-Kirby,A., Lajoie,G. and Heinrichs,D.E. (2004) Role of siderophore biosynthesis in virulence of Staphylococcus aureus: identification and characterization of genes involved in production of a siderophore. Infect. Immun., 72, 29–37.

84. Cheung,J., Beasley,F.C., Liu,S., Lajoie,G.A. and Heinrichs,D.E. (2009) Molecular characterization of staphyloferrin B biosynthesis in Staphylococcus aureus. Mol. Microbiol., 74, 594–608.

85. Repka,L.M., Chekan,J.R., Nair,S.K. and van der Donk,W.A. (2017) Mechanistic Understanding of Lanthipeptide Biosynthetic Enzymes. Chem. Rev., 117, 5457–5520.

86. Xu,M., Zhang,F., Cheng,Z., Bashiri,G., Wang,J., Hong,J., Wang,Y., Xu,L., Chen,X., Huang,S.-X., et al. (2020) Functional genome mining reveals a class V lanthipeptide containing a d-amino acid introduced by an F420 H2-dependent reductase. Angew. Chem. Int. Ed Engl., 59, 18029–18035.

87. Völler,G.H., Krawczyk,J.M., Pesic,A., Krawczyk,B., Nachtigall,J. and Süssmuth,R.D. (2012) Characterization of new class III lantibiotics--erythreapeptin, avermipeptin and griseopeptin from Saccharopolyspora erythraea, Streptomyces avermitilis and Streptomyces griseus demonstrates stepwise N-terminal leader processing. Chembiochem, 13, 1174–1183.

88. Ayikpoe,R., Govindarajan,V. and Latham,J.A. (2019) Occurrence, function, and biosynthesis of mycofactocin. Appl. Microbiol. Biotechnol., 103, 2903–2912.

89. Ellerhorst,M., Barth,S.A., Graça,A.P., Al-Jammal,W.K., Peña-Ortiz,L., Vilotijevic,I. and Lackner,G. (2022) S-Adenosylmethionine (SAM)-Dependent Methyltransferase MftM is Responsible for Methylation of the Redox Cofactor Mycofactocin. ACS Chem. Biol., 17, 3207–3217.

90. Hudson,G.A., Burkhart,B.J., DiCaprio,A.J., Schwalen,C.J., Kille,B., Pogorelov,T.V. and Mitchell,D.A. (2019) Bioinformatic mapping of radical S-adenosylmethionine-dependent ribosomally synthesized and post-translationally modified peptides identifies new Cα, Cβ, and Cγ-linked thioether-containing peptides. J. Am. Chem. Soc., 141, 8228–8238.

91. Peña-Ortiz,L., Graça,A.P., Guo,H., Braga,D., Köllner,T.G., Regestein,L., Beemelmanns,C. and Lackner,G. (2020) Structure elucidation of the redox cofactor mycofactocin reveals oligo-glycosylation by MftF. Chem. Sci., 11, 5182–5190.

92. Baltz,R.H. (2021) Genome mining for drug discovery: progress at the front end. J. Ind. Microbiol. Biotechnol., 48, kuab044.

93. Blin,K., Shaw,S., Kautsar,S.A., Medema,M.H. and Weber,T. (2021) The antiSMASH database version 3: increased taxonomic coverage and new query features for modular enzymes. Nucleic Acids Res., 49, D639–D643.

94. Ziemert,N., Lechner,A., Wietz,M., Millán-Aguiñaga,N., Chavarria,K.L. and Jensen,P.R. (2014) Diversity and evolution of secondary metabolism in the marine actinomycete genus Salinispora. Proc. Natl. Acad. Sci. U. S. A., 111, E1130–9.

95. Cimermancic,P., Medema,M.H., Claesen,J., Kurita,K., Wieland Brown,L.C., Mavrommatis,K., Pati,A., Godfrey,P.A., Koehrsen,M., Clardy,J., et al. (2014) Insights into secondary metabolism from a global analysis of prokaryotic biosynthetic gene clusters. Cell, 158, 412–421.

96. Doroghazi,J.R., Albright,J.C., Goering,A.W., Ju,K.-S., Haines,R.R., Tchalukov,K.A., Labeda,D.P., Kelleher,N.L. and Metcalf,W.W. (2014) A roadmap for natural product discovery based on large-scale genomics and metabolomics. Nat. Chem. Biol., 10, 963– 968.

97. Salamzade,R., Cheong,J.Z.A., Sandstrom,S., Swaney,M.H., Stubbendieck,R.M., Starr,N.L., Currie,C.R., Singh,A.M. and Kalan,L.R. (2023) Evolutionary investigations of the biosynthetic diversity in the skin microbiome using lsaBGC. Microbial Genomics, 9.

98. Steinegger,M. and Söding,J. (2018) Clustering huge protein sequence sets in linear time. Nat. Commun., 9, 2542.

99. Shen,W., Le,S., Li,Y. and Hu,F. (2016) SeqKit: A Cross-Platform and Ultrafast Toolkit for FASTA/Q File Manipulation. PLoS One, 11, e0163962.

100. Nguyen,L.-T., Schmidt,H.A., von Haeseler,A. and Minh,B.Q. (2015) IQ-TREE: a fast and effective stochastic algorithm for estimating maximum-likelihood phylogenies. Mol. Biol. Evol., 32, 268–274.

101. Hackl,T., Duponchel,S., Barenhoff,K., Weinmann,A. and Fischer,M.G. (2021) Virophages and retrotransposons colonize the genomes of a heterotrophic flagellate. Elife, 10.

102. Gilchrist,C.L.M. and Chooi,Y.-H. (2021) clinker & clustermap.js: automatic generation of gene cluster comparison figures. Bioinformatics, 37, 2473–2475.

103. Sievers,F., Wilm,A., Dineen,D., Gibson,T.J., Karplus,K., Li,W., Lopez,R., McWilliam,H., Remmert,M., Söding,J., et al. (2011) Fast, scalable generation of high-quality protein multiple sequence alignments using Clustal Omega. Mol. Syst. Biol., 7, 539.

104. Zhou,L., Feng,T., Xu,S., Gao,F., Lam,T.T., Wang,Q., Wu,T., Huang,H., Zhan,L., Li,L., et al. (2022) gams: a visual exploration tool for multiple sequence alignment and associated data. Brief. Bioinform., 23, 1–12.

