## Supplementary Information for "BGCFlow: Systematic pangenome workflow for the analysis of biosynthetic gene clusters across large genomic datasets"

**Table S1. List of 14 main computational tools to select the large-scale genome mining analysis using BGCFlow**

| | Rule Name | Tool Name | Version | Description | Link | Refs |
|---|---|---|---|---|---|---|
| 1 | eggnog | eggnog-mapper | 2.1.6 | Functional annotation of genome sequences using pre-computed Orthologous Group and phylogenies from the EggNOG database (http://eggnog5.embl.de). | https://github.com/eggnogdb/eggnog-mapper | (1, 2) |
| 2 | mash | mash | 2.3 | Calculate pairwise distance estimation for all samples using MinHash. | https://github.com/marbl/Mash | (3, 4) |
| 3 | fastani | fastani | 1.33 | Calculate pairwise Average Nucleotide Identity (ANI) across all samples. | https://github.com/ParBLiSS/FastANI | (5) |
| 4 | automlst_wrapper | Automlst-wapper | 2bccf68 | Simplified Species Tree building of all samples using [autoMLST](https://github.com/NBChub/automlst-simplified-wrapper) | https://github.com/KatSteinke/automlst-simplified-wrapper | (6) |
| 5 | roary | roary | 3.13.0 | Build pangenome from all samples using Roary. | https://github.com/sanger-pathogens/Roary | (7) |
| 6 | seqfu_stats | Seqfu2 | 1.15.3 | Calculate sequence statistics using SeqFu. | https://github.com/telatin/seqfu2 | (8) |
| 7 | bigslice | bigslice | ba4056a | Cluster BGCs using BiG-SLiCE (https://github.com/medema-group/bigslice) or map BGCs to BiG-FAM database (https://bigfam.bioinformatics.nl/) | https://github.com/medema-group/bigslice | (9, 10) |
| 8 | checkm | checkm-genome | 1.1.3 | Assess genome quality with CheckM. | https://github.com/Ecogenomics/CheckM | (11) |
| 9 | gtdbtk | gtdbtk | 2.1.0 | Taxonomic placement with GTDB-Tk with GTDB release 207 | https://github.com/Ecogenomics/GTDBTk | (12, 13) |
| 10 | prokka | prokka | 1.14.6 | Copy annotated genbank results. | https://github.com/tseemann/prokka | (14) |
| 11 | antismash | antismash | 6.1.1 | Summarizes antiSMASH result. | https://github.com/antismash | (15) |
| 12 | arts | arts | 3e32474 | Targeted genome mining with Antibiotic Resistant Target Seeker (ARTS2) on samples. | https://github.com/NBChub/arts_v3 | (16, 17) |
| 13 | deeptfactor | DeepTFactor | 7f1bcb4 | Use deep learning to find Transcription Factors. | https://bitbucket.org/kaistsystemsbiology/deeptfactor/src | (18) |
| 14 | bigscape | BiG-SCAPE | 1.1.4 | Cluster BGCs using BiG-SCAPE | https://github.com/medema-group/BiG-SCAPE | (19) |

### Table S2. Rule names in BGCFlow main Snakemake workflow

| node_number | Rule Name | Description | Refs |
|---|---|---|---|
| 1 | fix_gtdb_taxonomy | Gather and fix taxonomy metadata into a summary table | |
| 2 | gtdb_prep | Fetch taxonomic information from publicly available genomes using GTDB API | (20) |
| 3 | seqfu_stats | Calculate sequence statistics using SeqFu. | (8) |
| 4 | mash | Calculate pairwise distance estimation for all samples using MinHash. | (3, 4) |
| 5 | fastani | Calculate pairwise Average Nucleotide Identity (ANI) across all samples. | (5) |
| 6 | checkm | Assess genome quality with CheckM. | (11) |
| 7 | install_checkm | Install CheckM locally | |
| 8 | install_gtdbtk | Install GTDB-tk locally | |
| 9 | prepare_gtdbtk_input | Prepare input files for GTDB-tk | |
| 10 | format_gbk | Correct naming and add taxonomy metadata in annotated genbank files | |
| 11 | prokka | Annotate bacterial genomes with prodigal and Prokka | (14) |
| 12 | extract_meta_prokka | Extract any taxonomy information from fasta files |  |
| 13 | bgc_count | Summarizes BGC count of a given antiSMASH result | |
| 14 | antismash | Detection of Biosynthetic Gene Clusters with antiSMASH | (15) |
| 15 | antismash_db_setup | Set up databases required for antiSMASH locally | |
| 16 | copy_antismash | Generate symlinks of antiSMASH result in the processed folder | |
| 17 | antismash_overview_gather | Compile all antiSMASH summary pages into a table | |
| 18 | antismash_overview | Extract the antiSMASH summary page of a given antiSMASH result | |
| 19 | downstream_bgc_prep | Prepare region genbank files and metadata for downstream analysis | |
| 20 | query_bigslice | Map BGCs to BiG-FAM database (https://bigfam.bioinformatics.nl/) | (9, 10) |
| 21 | fetch_bigslice_db | Install BiG-FAM database locally | |
| 22 | bigscape | Cluster BGCs using BiG-SCAPE | (19) |
| 23 | install_bigscape | Install BiG-SCAPE locally | |
| 24 | bigscape_to_cytoscape | Generate cytoscape ready tables and annotation files | |
| 25 | get_mibig_table | Get MIBIG 3.0 tables locally | (21) |
| 26 | bigslice_prep | Prepare files and metadata required for BiG-SLICE clustering | |
| 27 | automlst_wrapper | Simplified Species Tree building of all samples using [autoMLST](https://github.com/NBChub/automlst-simplified-wrapper) | (6) |
| 28 | prep_automlst_gbk | Prepare input files for autoMLST | |
| 29 | install_automlst_wrapper | Install autoMLST locally | |

| node_number | Rule Name | Description | Refs |
|---|---|---|---|
| 30 | arts_extract | Extract BGC proximity hits summary from an ARTS result | |
| 31 | arts | Targeted genome mining with Antibiotic Resistant Target Seeker (ARTS2) on samples. | (16, 17) |
| 32 | install_eggnog | Install Eggnog databases locally | |
| 33 | deeptfactor_to_json | Convert DeepTFactor result into json table | |
| 34 | deeptfactor | Use deep learning to find Transcription Factors. | (18) |
| 35 | deeptfactor_setup | Install DeepTFactor locally | |
| 36 | roary | Build pangenome from all samples using Roary. | (7) |
| 37 | ncbi_genome_download | Download NCBI genome assemblies from RefSeq or GenBank using ncbi-genome-download (https://github.com/kblin/ncbi-genome-download) | |
| 38 | patric_genome_download | Fetch genome fasta files from Patric database | |
| 39 | copy_custom_fasta | Grab user-provided input fasta file for processing | |
| 40 | seqfu_combine | Combine all seqfu result into a table | |
| 41 | checkm_out | Extract and summarizes CheckM results | |
| 42 | extract_ncbi_information | Capture metadata of downloaded genomes from NCBI | |
| 43 | copy_prokka_gbk | Get a copy of annotated genbank files into processed folder | |
| 44 | eggnog | Functional annotation of genome sequences using pre-computed Orthologous Group and phylogenies from the EggNOG database (http://eggnog5.embl.de). | (1, 2) |
| 45 | deeptfactor_summary | Combine all DeepTFactor result into a table | |
| 46 | mash_convert | Convert MASH distance result into pandas ready matrix | |
| 47 | fastani_convert | Convert FastANI result into pandas ready matrix | |
| 48 | gtdbtk | Taxonomic placement with GTDB-Tk with GTDB release 207 | (12, 13) |
| 49 | automlst_wrapper_out | Extract and format autoMLST tree into processed folder | |
| 50 | antismash_summary | Correct shortened accession and compile antiSMASH summary into a table | |
| 51 | arts_combine | Combine ARTS result into a table | |
| 52 | cblaster_genome_db | Build diamond database of genomes for cblaster search. | (22, 23) |
| 53 | summarize_bigslice_query | Summarize BiG-FAM GCF hits | |
| 54 | copy_bigscape | Copy and format BiG-SCAPE result into processed folder | |
| 55 | bigslice | Cluster BGCs using BiG-SLiCE (https://github.com/medema-group/bigslice) | (10) |
| 56 | cblaster_bgc_db | Build diamond database of BGCs for cblaster search. | (22, 23) |

| node_number | Rule Name | Description | Refs |
|---|---|---|---|
| 57 | roary_out | Extract ROARY information | |
| 58 | eggnog_roary | Functional annotation of Roary output using eggNOG mapper | (1, 2, 7) |
| 59 | deeptfactor_roary | Use DeepTFactor on Roary outputs. | (7, 18) |

a)

```yaml
# This file should contain everything to configure the workflow on a global scale.

#### PROJECT INFORMATION ####
# This section control your project configuration.
# Each project are separated by "-".
# A project must contain the variable "name" and "samples".
# - name : name of your project
# - samples : a csv file containing a list of genome ids for analysis with multiple sources mentioned. Genome ids must be unique.
# - prokka-db (optional): list of the custom accessions to use as prokka reference database
# - gtdb-tax (optional): output summary file of GTDB-tk with "user_genome" and "classification" as the two minimum columns

projects:
# Project 1
- name: config/qc_saccharopolyspora/project_config.yaml

# Project 2
 - name: config/mq_saccharopolyspora/project_config.yaml

#### RESOURCES CONFIGURATION ####
# resources : the location of the resources to run the rule.
# The default location is at "resources/{resource_name}".
resources_path:
  antismash_db: resources/antismash_db
  eggnog_db: resources/eggnog_db
  BiG-SCAPE: resources/BiG-SCAPE
  bigslice: resources/bigslice
  checkm: resources/checkm
  #RNAmmer: resources/RNAmmer # Path must be specified if rule configuration "rnammer" is TRUE
```

b)

```yaml
name: mq_saccharopolyspora
pep_version: 2.1.0
description: '28 Saccharopolysopra genomes of medium to high quality with less than 60 contigs'
sample_table: samples.csv
prokka-db: prokka-db.csv
gtdb-tax: 'gtdbtk.bac120.summary.tsv'

rules:
  seqfu: TRUE
  mash: TRUE
  fastani: TRUE
  checkm: FALSE
  gtdbtk: FALSE
  prokka-gbk: FALSE
  antismash: TRUE
  query-bigslice: TRUE
  bigscape: TRUE
  bigslice: FALSE
  automlst-wrapper: TRUE
  arts: TRUE
  roary: FALSE
  eggnog: FALSE
  eggnog-roary: FALSE
  deeptfactor: FALSE
  deeptfactor-roary: FALSE
  cblaster-genome: FALSE
  cblaster-bgc: FALSE
```

c)

| 1 | genome_id | source | organism | genus | species | strain |
|---|---|---|---|---|---|---|
| 2 | GCF_003635025.1 | ncbi | | | | |
| 3 | GCF_900114905.1 | ncbi | | | | |
| 4 | GCF_007829955.1 | ncbi | | | | |
| 5 | GCF_018070075.1 | ncbi | | | | |
| 6 | GCF_016859185.1 | ncbi | | | | |
| 7 | GCF_018141105.1 | ncbi | | | | |
| 8 | GCF_022392385.1 | ncbi | | | | |
| 9 | GCF_000062885.1 | ncbi | | | | |
| 10 | GCF_002564065.1 | ncbi | | | | |
| 11 | GCF_900116135.1 | ncbi | | | | |
| 12 | GCF_014203325.1 | ncbi | | | | |
| 13 | GCF_022828475.1 | ncbi | | | | |
| 14 | GCF_024734405.1 | ncbi | | | | |
| 15 | GCF_008630535.1 | ncbi | | | | |
| 16 | GCF_013410345.1 | ncbi | | | | |
| 17 | GCF_900112555.1 | ncbi | | | | |
| 18 | GCF_900108315.1 | ncbi | | | | |
| 19 | GCF_014203395.1 | ncbi | | | | |
| 20 | GCF_000710755.1 | ncbi | | | | |
| 21 | GCF_003931915.1 | ncbi | | | | |
| 22 | GCF_025643595.1 | ncbi | | | | |
| 23 | GCF_012277335.1 | ncbi | | | | |
| 24 | GCF_015209505.1 | ncbi | | | | |
| 25 | GCF_016526145.1 | ncbi | | | | |
| 26 | GCF_014490055.1 | ncbi | | | | |
| 27 | GCF_002846475.1 | ncbi | | | | |
| 28 | GCF_000194155.1 | ncbi | | | | |
| 29 | GCF_014646075.1 | ncbi | | | | |

d)

| 1 | Accession | Strain Description |
|---|---|---|
| 2 | GCA_000062885.1 | Saccharopolyspora erythraea NRRL 2338 (high GC Gram+) |
| 3 | GCA_000009765.2 | Streptomyces avermitilis MA-4680 = NBRC 14893 (high GC Gram+) |
| 4 | GCA_000010605.1 | Streptomyces griseus subsp. griseus NBRC 13350 (high GC Gram+) |
| 5 | GCA_000015005.1 | Mycolicibacterium smegmatis MC2 155 (high GC Gram+) |
| 6 | GCA_000196835.1 | Amycolatopsis mediterranei U32 (high GC Gram+) |
| 7 | GCA_000203835.1 | Streptomyces coelicolor A3(2) (high GC Gram+) |
| 8 | GCA_000444875.1 | Streptomyces collinus Tu 365 (high GC Gram+) |
| 9 | GCA_000739105.1 | Streptomyces lividans TK24 (high GC Gram+) |

**Figure S1. Project configuration and metadata to setup BGCFlow**

a) Global configuration file (*yaml*) with path to two of the PEPs used in the demonstration. b) Example of the *mq_saccharopolyspora* PEP configuration file with selected rules to run. c) List of 28 NCBI accession IDs in the samples table used in the project (*samples.csv*). d) List of selected high-quality genome annotations of actinomycetes as priority of prokka annotation. (*prokka-db.csv*)

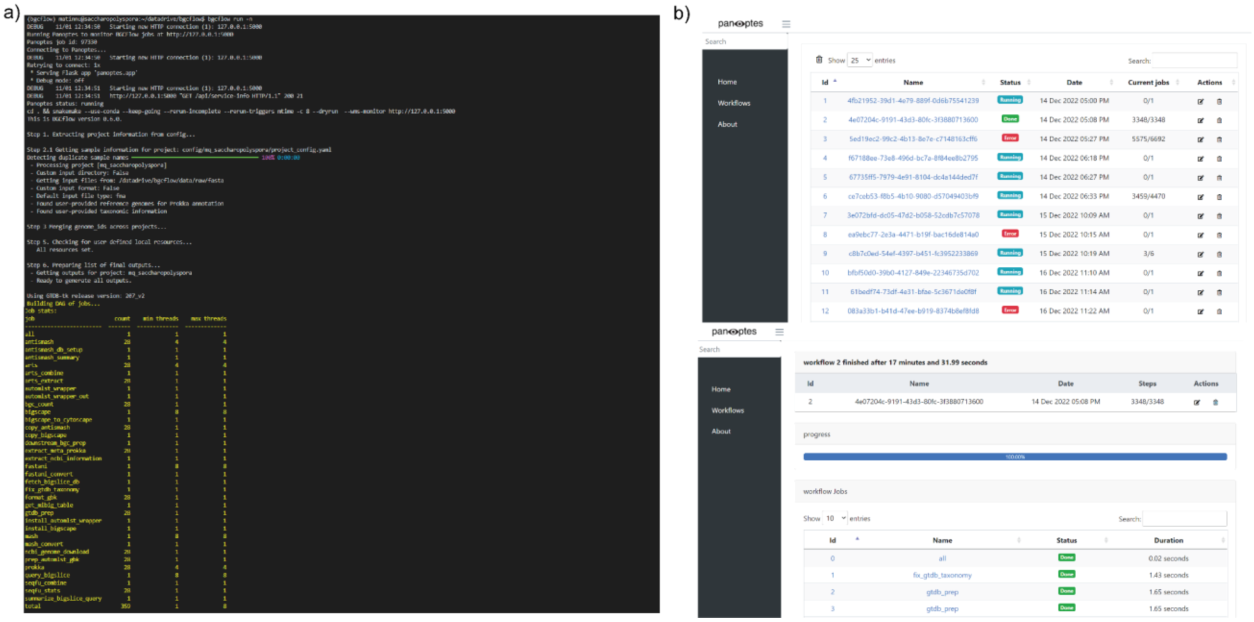

**Figure S2. Job scheduling and monitoring using Snakemake and panoptes**

a) Screenshot of the dry run for BGCFlow on the *mq_saccharopolyspora* with all jobs to run for the selected rules in Figure S2b. The number of minimum thread for each jobs can be updated using configure file for the thread profiling. b) Monitoring of the Snakemake runs and individual jobs using panoptes.

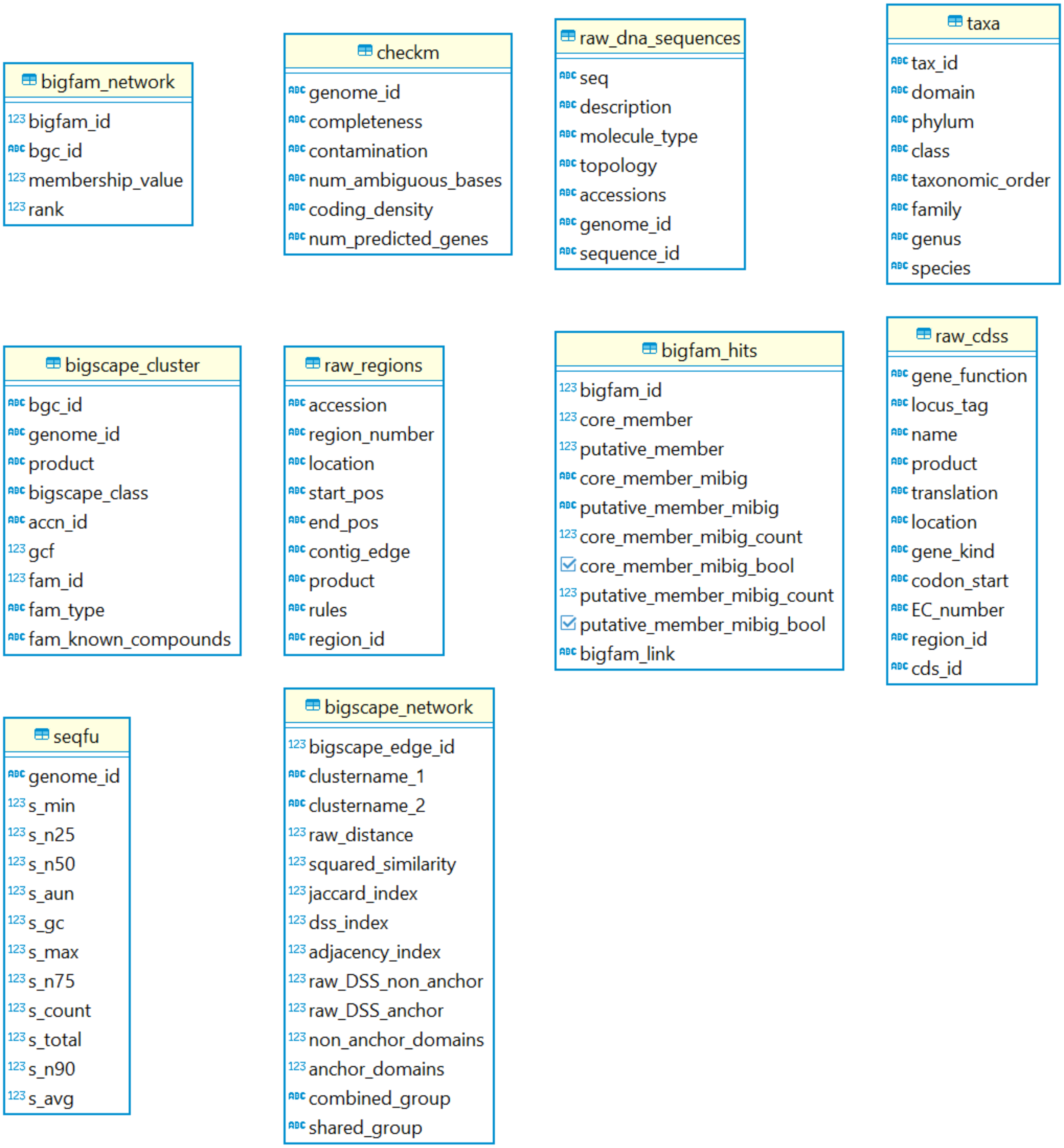

**Figure S3. Entity Relations Diagram of the DuckDB OLAP database**

Entity relations diagram of the exported DuckDB tables using DBT. Schema are being maintained in https://github.com/NBChub/bgcflow_dbt-duckdb.

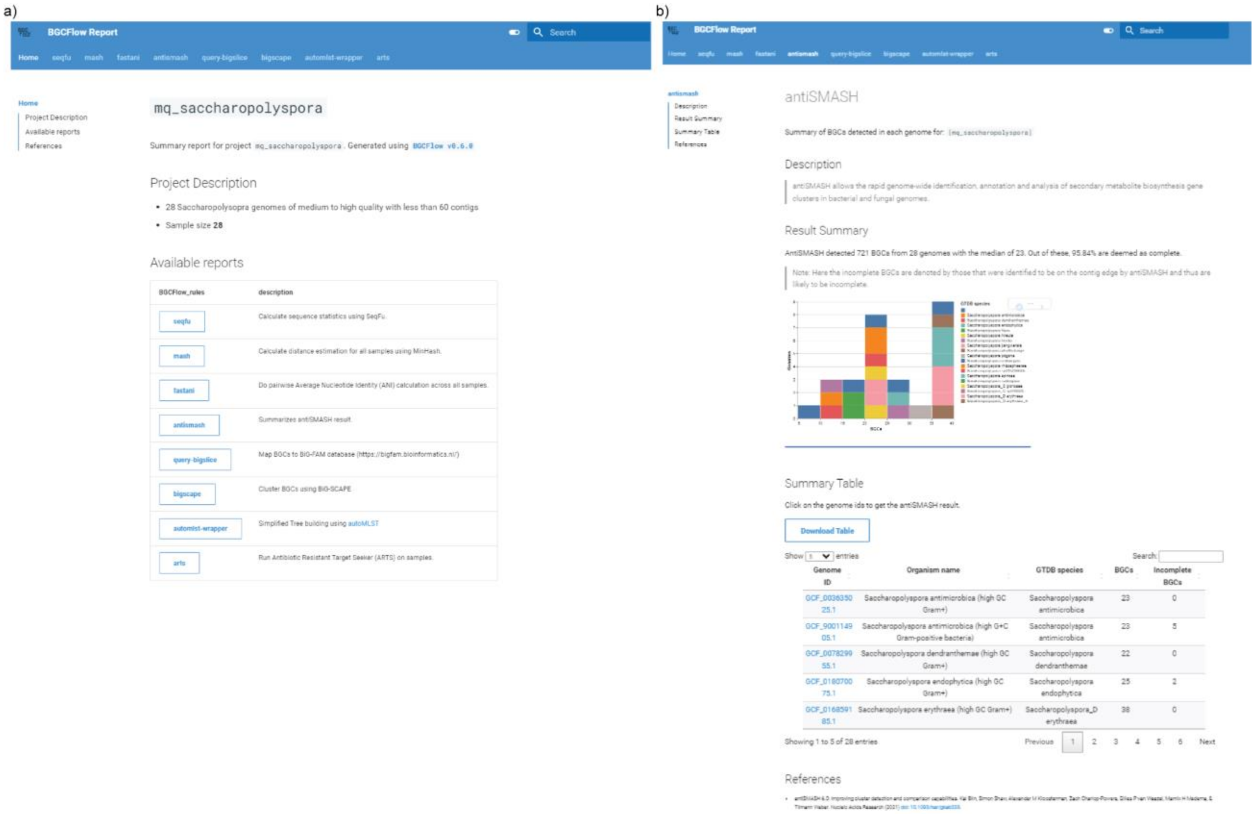

**Figure S4. Example of Jupyter-based markdown reports**

a) Screenshot of the home page of the BGCFlow reports of the *mq_saccharopolyspora* project with selected rules. b) Example report of the *antismash* rule's report page with a summary and the table with a list of genomes. The links for the genomes IDs in the table correspond to antiSMASH-generated HTML reports with details on all detected BGCs.

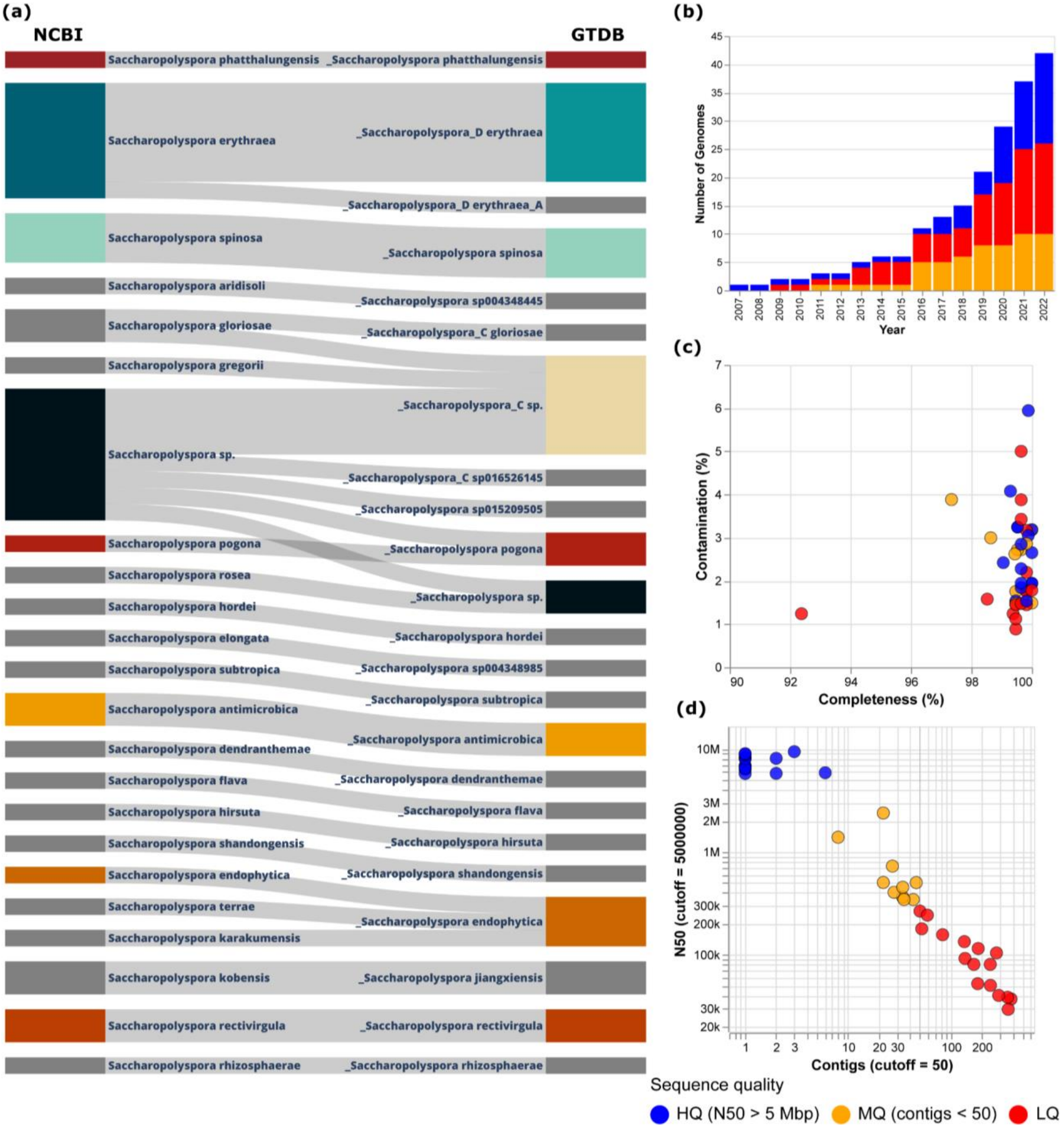

**Figure S5. Overview of timeline, quality, and taxonomic placement of 42 *Saccharopolyspora* genomes**

a) Cumulative bar chart of the number of genomes over the last 15 years with different assembly qualities. b) Distribution of contamination vs completeness metrics calculated using CheckM, where colors represent the assembly qualities. c) Sankey diagram representing the species assignment differences between NCBI and GTDB. d) Scatterplot representing the distribution of N50 values vs the number of contigs. The cutoff of 50 contigs is used to filter the low-quality genomes, whereas 5 Mbp of N50 value cutoff was used to define high-quality genomes. The remaining genomes were defined as medium-quality.

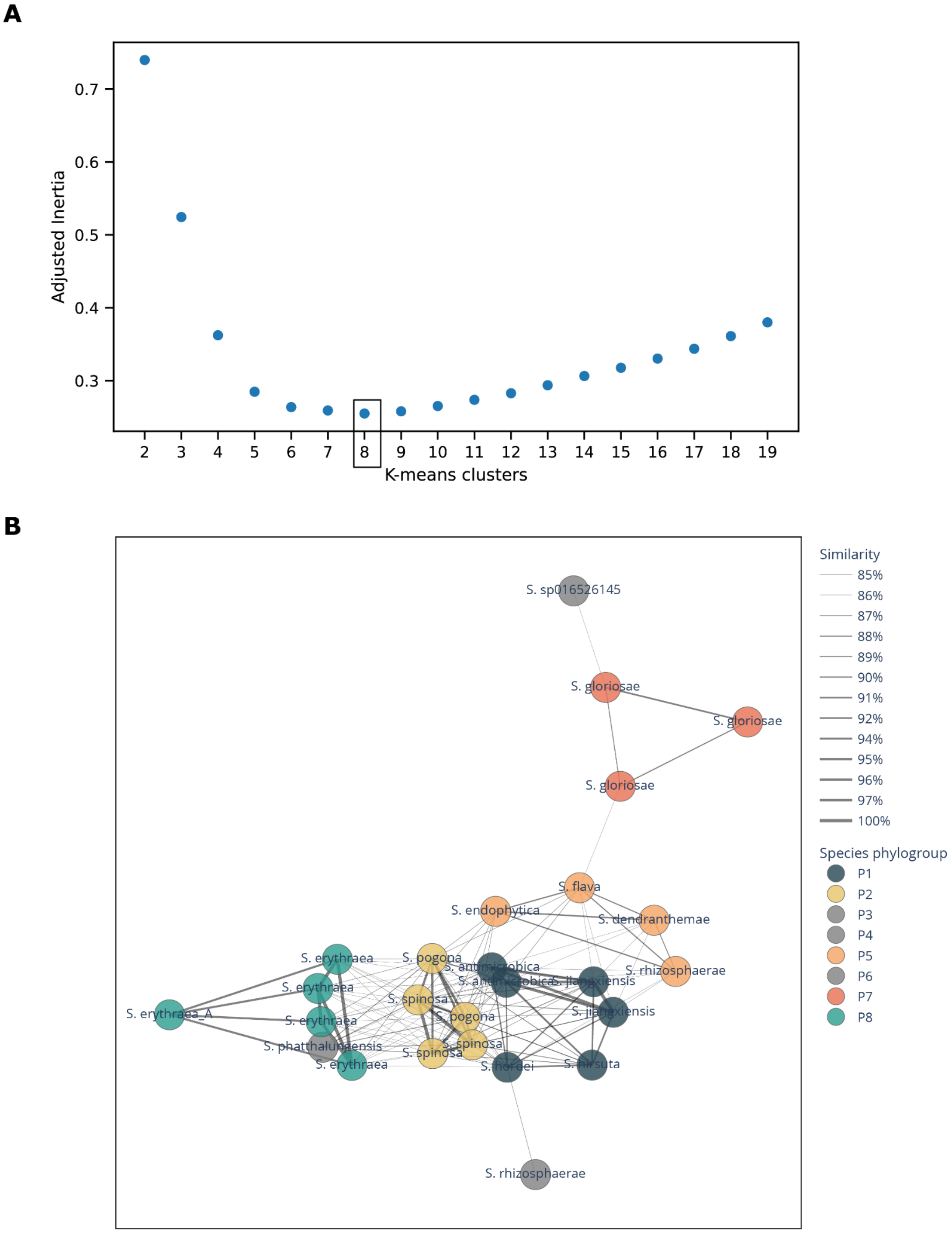

**Figure S6. Report of genome clustering using Mash distances.**
(a) Suggested optimal clustering based on K-means clustering with minimum adjusted inertia. (b) MASH-similarity network with nodes represented by genomes from different species and edge width represented by the similarity of 85% to 100%. The 8 optimal clusters are represented by different colors of nodes.

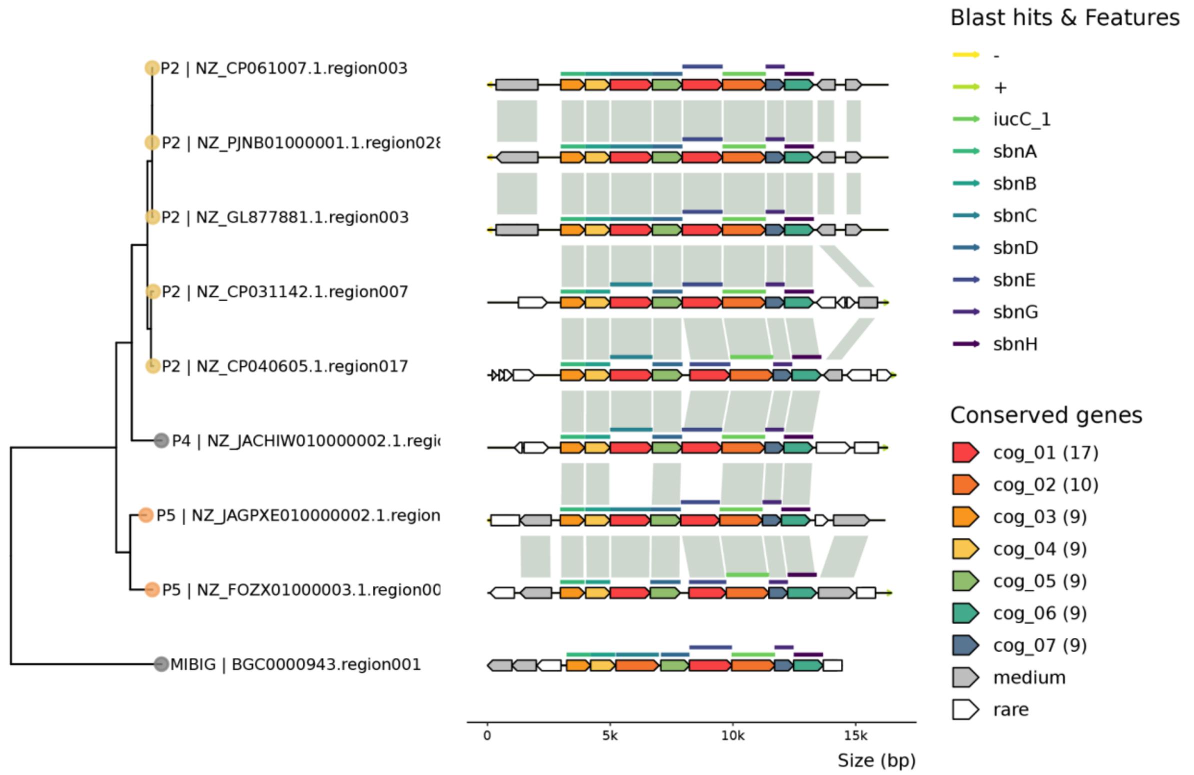

**Figure S7. Gene comparison of staphylobactin-like BGCs in *Saccharopolyspora***
Comparison of staphylobactin-like BGCs which are connected through a BiG-FAM node *GCF_201888,* which has a Shannon index of ~0.3 and contained 12,444 BGCs which are distributed across 43 genera with the majority belonging to Staphylococcus (~94.2%). All 8 genes responsible for staphylobactin biosynthesis can be mapped through CBlaster except for NZ_FOZX01000003, with only 7 matches.

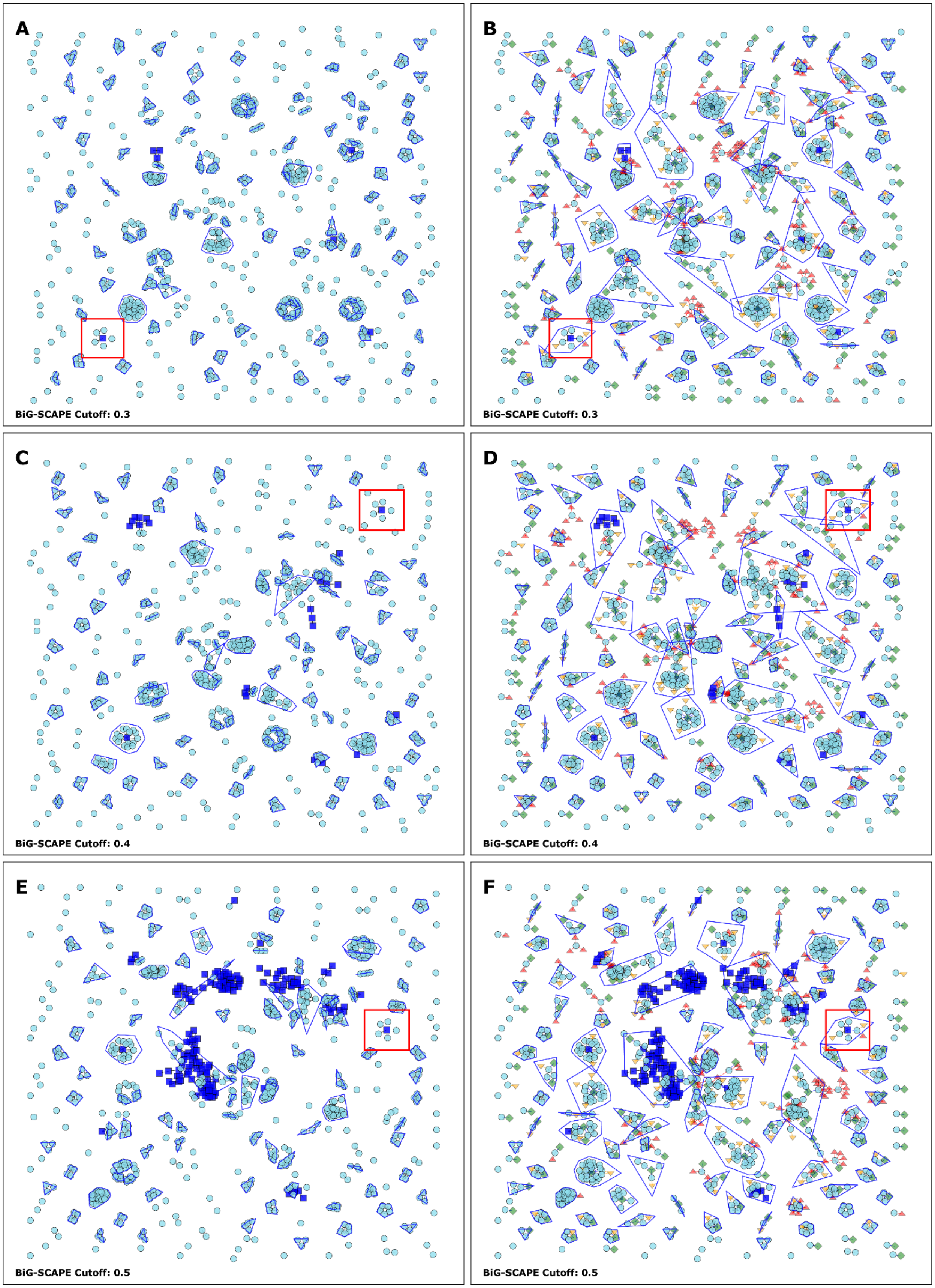
A
BiG-SCAPE Cutoff: 0.3
B
BiG-SCAPE Cutoff: 0.3
C
BiG-SCAPE Cutoff: 0.4
D
BiG-SCAPE Cutoff: 0.4
E
BiG-SCAPE Cutoff: 0.5
F
BiG-SCAPE Cutoff: 0.5

47

**Figure S8. Comparison of BiG-SCAPE network with different cutoffs assignment with the enriched network**

This figure compares the number of connected component assignments between an enriched and non-enriched BiG-SCAPE network with different cutoffs. BiG-SCAPE networks were enriched with ARTS2 hits, BiG-FAM hits, and KnownClusterBlast hits. BiG-FAM models 202087, 200946, 210179, 213140, 201682, 201608, 205957, 215277, 201830, and 202082 were removed because the assigned top genus in the model is below 30%. A) BiG-SCAPE sequence similarity network with cutoff 0.3 resulting in 328 connected components (GCFs) with 206 singletons. A total of 4 GCFs with 14 BGCs can be assigned to 6 MIBIG entries. B) Enriched BiG-SCAPE network with cutoff 0.3 resulting in 202 connected components with 44 singletons. This increases the number of BGCs in the 4 BiG-SCAPE GCF with MIBIG hits into 29 BGCs. Additionally, a total of 105 BGCs can be assigned to 12 GCFs with MIBIG KnownClusterBlast similarity >= 80% and left only 122 BGCs in 70 GCFs without connection to MIBIG and BiG-FAM nodes. C) BiG-SCAPE sequence similarity network with cutoff 0.4 resulting in 280 connected components (GCFs) with 162 singletons. A total of 10 GCFs with 80 BGCs can be assigned to 19 MIBIG entries. D) Enriched BiG-SCAPE network with cutoff 0.4 resulting in 190 connected components with 41 singletons. This increases the number of BGCs in the 10 BiG-SCAPE GCF with MIBIG hits into 109 BGCs. Additionally, a total of 53 BGCs can be assigned to 8 GCFs with MIBIG KnownClusterBlast similarity >= 80% and left only 115 BGCs in 64 GCFs without connection to MIBIG and BiG-FAM nodes. E) BiG-SCAPE sequence similarity network with cutoff 0.5 resulting in 259 connected components (GCFs) with 144 singletons. A total of 15 GCFs with 115 BGCs can be assigned to 110 MIBIG entries. F) Enriched BiG-SCAPE network with cutoff 0.5 resulting in 173 connected components with 36 singletons. This increases the number of BGCs in the 15 BiG-SCAPE GCF with MIBIG hits into 184 BGCs. Additionally, a total of 44 BGCs can be assigned to 7 GCFs with MIBIG KnownClusterBlast similarity >= 80% and left only 114 BGCs in 59 GCFs without connection to MIBIG and BiG-FAM nodes. Red box highlights the location of spinosyn-like BGCs in the network.

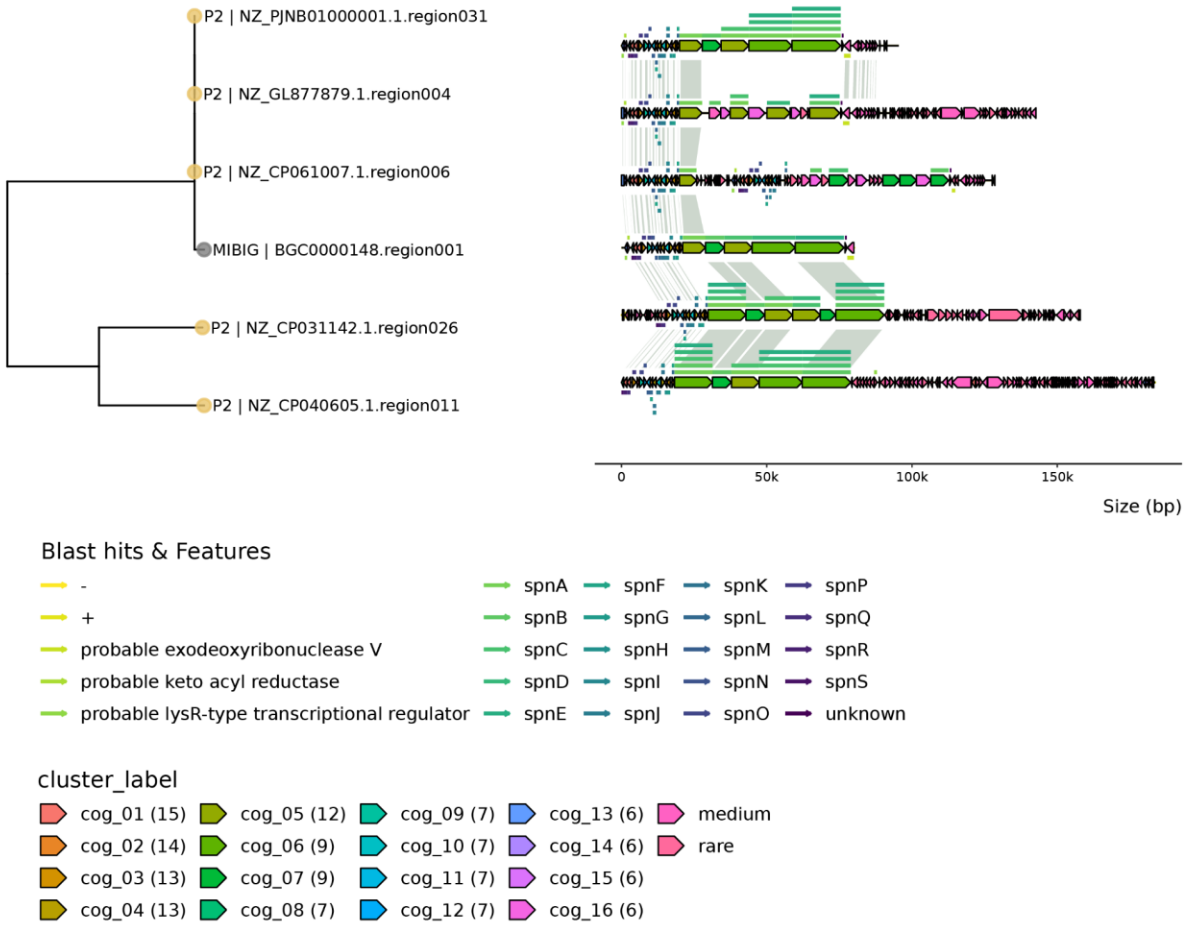

**Figure S9. Gene comparison of spinosyn-like BGCs in *Saccharopolyspora***

Comparison of BGCs with BiG-SCAPE and KnownClusterBlast similarity to spinosyn. All regions matched with spinosyn belong to phylogroup 2. Of the 5 BGCs, only 1 has an exact structure with spinosyn MIBIG BGC (NZ_PJNB01000001). Other BGCs showed variation in the PKS modules and might even lose the whole biosynthesis pathway (NZ_CP061007). Annotations in all BGCs showed matches to specific tailoring enzymes found in spinosyn biosynthesis for sugar attachment (cog_03 - probable rhamonsyltransferase).

**Data S1. Input and result tables related to the PEP on qc_saccharaopolyspora**

Tab 1. Content of the sample.csv listing 42 NCBI genomes as input for the BGCFlow PEP configuration
Tab 2. NCBI metadata of the 42 NCBI genomes
Tab 3. CheckM results on quality assessment of the 42 NCBI genomes
Tab 4. SeqFu results on quality assessment of the 42 NCBI genomes
Tab 5. GTDB-tk and GTDB results on taxonomic definition of the 42 NCBI genomes

**Data S2. Input and result tables related to the PEP on mq_saccharaopolyspora**

Tab 1. Content of the sample.csv listing 26 genomes as input for the BGCFlow PEP configuration
Tab 2. MASH distance of the selected 26 genomes
Tab 3. MASH-base phlyogroups of the selected 26 genomes
Tab 4. Prokka annotation result summary of the selected 26 genomes
Tab 5. List of BGC hits from antiSMASH
Tab 6. Results of GCFs based on BiG-SCAPE using 0.3 cutoff
Tab 7. Results of GCFs based on BiG-SCAPE using 0.4 cutoff
Tab 8. Results of GCFs based on BiG-SCAPE using 0.5 cutoff
Tab 9. Edge table of BiG-SCAPE sequence similarity network using 0.3 cutoff
Tab 10. Edge table of BiG-SCAPE sequence similarity network using 0.4 cutoff
Tab 11. Edge table of BiG-SCAPE sequence similarity network using 0.5 cutoff

**Data S3. Results of BiG-FAM and ARTS database related to the PEP on mq_saccharaopolyspora**

Tab 1. Hits against the BiG-FAM GCFs calculated using BiG-SLICE query
Tab 2. List of BiG-FAM GCFs hits
Tab 3. Hits against ARTS profile
Tab 4. Table with all edges represented in the enriched network with BiG-SCAPE cutoff 0.3

**Data S4. Results of detailed GCF comparison related to the PEP on staphylobactin-like BGCs**

Tab 1. CDS Feature and COG annotations of staphylobactin-like GCF
Tab 2. Clinker links for BGC alignment of staphylobactin-like GCF
Tab 3. CBlaster hits for BGC alignment of staphylobactin-like GCF

**Data S5. Results of detailed GCF comparison related to the PEP on spinosyn-like BGCs**

Tab 1. CDS Feature and COG annotations of spinosyn-like GCF
Tab 2. Clinker links for BGC alignment of spinosyn-like GCF
Tab 3. CBlaster hits for BGC alignment of spinosyn-like GCF

**Data S6. Results of detailed GCF comparison related to the PEP on erythreapeptin-like BGCs**

Tab 1. CDS Feature and COG annotations of erythreapeptin-like GCF
Tab 2. Clinker links for BGC alignment of erythreapeptin-like GCF
Tab 3. CBlaster hits for BGC alignment of erythreapeptin-like GCF

**Data S7. Results of detailed GCF comparison related to the PEP on mycofactocin-like BGCs**

Tab 1. CDS Feature and COG annotations of mycofactocin-like GCF
Tab 2. Clinker links for BGC alignment of mycofactocin-like GCF
Tab 3. CBlaster hits for BGC alignment of mycofactocin-like GCF

### References

1. Cantalapiedra,C.P., Hernández-Plaza,A., Letunic,I., Bork,P. and Huerta-Cepas,J. (2021) eggNOG-mapper v2: Functional Annotation, Orthology Assignments, and Domain Prediction at the Metagenomic Scale. *Mol. Biol. Evol.*, **38**, 5825–5829.

2. Huerta-Cepas,J., Szklarczyk,D., Heller,D., Hernández-Plaza,A., Forslund,S.K., Cook,H., Mende,D.R., Letunic,I., Rattei,T., Jensen,L.J., *et al.* (2019) eggNOG 5.0: a hierarchical, functionally and phylogenetically annotated orthology resource based on 5090 organisms and 2502 viruses. *Nucleic Acids Res.*, **47**, D309–D314.

3. Ondov,B.D., Treangen,T.J., Melsted,P., Mallonee,A.B., Bergman,N.H., Koren,S. and Phillippy,A.M. (2016) Mash: fast genome and metagenome distance estimation using MinHash. *Genome Biol.*, **17**, 132.

4. Ondov,B.D., Starrett,G.J., Sappington,A., Kostic,A., Koren,S., Buck,C.B. and Phillippy,A.M. (2019) Mash Screen: high-throughput sequence containment estimation for genome discovery. *Genome Biol.*, **20**, 232.

5. Jain,C., Rodriguez-R,L.M., Phillippy,A.M., Konstantinidis,K.T. and Aluru,S. (2018) High throughput ANI analysis of 90K prokaryotic genomes reveals clear species boundaries. *Nat. Commun.*, **9**, 5114.

6. Alanjary,M., Steinke,K. and Ziemert,N. (2019) AutoMLST: an automated web server for generating multi-locus species trees highlighting natural product potential. *Nucleic Acids Res.*, **47**, W276–W282.

7. Page,A.J., Cummins,C.A., Hunt,M., Wong,V.K., Reuter,S., Holden,M.T.G., Fookes,M., Falush,D., Keane,J.A. and Parkhill,J. (2015) Roary: rapid large-scale prokaryote pan genome analysis. *Bioinformatics*, **31**, 3691–3693.

8. Telatin,A., Fariselli,P. and Birolo,G. (2021) SeqFu: A Suite of Utilities for the Robust and Reproducible Manipulation of Sequence Files. *Bioengineering (Basel)*, **8**.

9. Kautsar,S.A., Blin,K., Shaw,S., Weber,T. and Medema,M.H. (2021) BiG-FAM: the biosynthetic gene cluster families database. *Nucleic Acids Res.*, **49**, D490–D497.

10. Kautsar,S.A., van der Hooft,J.J.J., de Ridder,D. and Medema,M.H. (2021) BiG-SLiCE: A highly scalable tool maps the diversity of 1.2 million biosynthetic gene clusters. *Gigascience*, **10**.

11. Parks,D.H., Imelfort,M., Skennerton,C.T., Hugenholtz,P. and Tyson,G.W. (2015) CheckM: assessing the quality of microbial genomes recovered from isolates, single cells, and metagenomes. *Genome Res.*, **25**, 1043–1055.

12. Chaumeil,P.-A., Mussig,A.J., Hugenholtz,P. and Parks,D.H. (2019) GTDB-Tk: a toolkit to classify genomes with the Genome Taxonomy Database. *Bioinformatics*, **36**, 1925–1927.

13. Parks,D.H., Chuvochina,M., Waite,D.W., Rinke,C., Skarshewski,A., Chaumeil,P.-A. and Hugenholtz,P. (2018) A standardized bacterial taxonomy based on genome phylogeny substantially revises the tree of life. *Nat. Biotechnol.*, **36**, 996–1004.

14. Seemann,T. (2014) Prokka: rapid prokaryotic genome annotation. *Bioinformatics*, **30**, 2068–2069.

15. Blin,K., Shaw,S., Kloosterman,A.M., Charlop-Powers,Z., van Wezel,G.P., Medema,M.H. and Weber,T. (2021) antiSMASH 6.0: improving cluster detection and comparison capabilities. *Nucleic Acids Res.*, **49**, W29–W35.

16. Mungan,M.D., Alanjary,M., Blin,K., Weber,T., Medema,M.H. and Ziemert,N. (2020) ARTS 2.0: feature updates and expansion of the Antibiotic Resistant Target Seeker for comparative genome mining. *Nucleic Acids Res.*, **48**, W546–W552.

17. Alanjary,M., Kronmiller,B., Adamek,M., Blin,K., Weber,T., Huson,D., Philmus,B. and Ziemert,N. (2017) The Antibiotic Resistant Target Seeker (ARTS), an exploration engine for antibiotic cluster prioritization and novel drug target discovery. *Nucleic Acids Res.*, **45**, W42–W48.

18. Kim,G.B., Gao,Y., Palsson,B.O. and Lee,S.Y. (2021) DeepTFactor: A deep learning-based tool for the prediction of transcription factors. *Proc. Natl. Acad. Sci. U. S. A.*, **118**.

19. Navarro-Muñoz,J.C., Selem-Mojica,N., Mullowney,M.W., Kautsar,S.A., Tryon,J.H., Parkinson,E.I., De Los Santos,E.L.C., Yeong,M., Cruz-Morales,P., Abubucker,S., *et al.* (2020) A computational framework to explore large-scale biosynthetic diversity. *Nat. Chem. Biol.*, **16**, 60–68.

20. Parks,D.H., Chuvochina,M., Rinke,C., Mussig,A.J., Chaumeil,P.-A. and Hugenholtz,P. (2022) GTDB: an ongoing census of bacterial and archaeal diversity through a phylogenetically consistent, rank normalized and complete genome-based taxonomy. *Nucleic Acids Res.*, **50**, D785–D794.

21. Terlouw,B.R., Blin,K., Navarro-Muñoz,J.C., Avalon,N.E., Chevrette,M.G., Egbert,S., Lee,S., Meijer,D., Recchia,M.J.J., Reitz,Z.L., *et al.* (2022) MIBiG 3.0: a community-driven effort to annotate experimentally validated biosynthetic gene clusters. *Nucleic Acids Res.*, **51**, gkac1049.

22. Buchfink,B., Reuter,K. and Drost,H.-G. (2021) Sensitive protein alignments at tree-of-life scale using DIAMOND. *Nat. Methods*, **18**, 366–368.

23. Gilchrist,C.L.M., Booth,T.J., van Wersch,B., van Grieken,L., Medema,M.H. and Chooi,Y.-H. (2021) cblaster: a remote search tool for rapid identification and visualization of homologous gene clusters. *Bioinformatics Advances*, **1**, vbab016.
